# Multiple-testing corrections in selection scans using identity-by-descent segments

**DOI:** 10.1101/2025.01.29.635528

**Authors:** Seth D. Temple, Sharon R. Browning

## Abstract

Failing to correct for multiple testing in selection scans can lead to false discoveries of recent genetic adaptations. The scanning statistics in selection studies are often too complicated to theoretically derive a genome-wide significance level or empirically validate control of the family-wise error rate (FWER). By modeling the autocorrelation of identity-by-descent (IBD) rates, we propose a computationally efficient method to determine genome-wide significance levels in an IBD-based scan for recent positive selection. In whole genome simulations, we show that our method has approximate control of the FWER and can adapt to the spacing of tests along the genome. We also show that these scans can have more than fifty percent power to reject the null model in hard sweeps with a selection coefficient *s >*= 0.01 and a sweeping allele frequency between twenty-five and seventy-five percent. A few human genes and gene complexes have statistically significant excesses of IBD segments in thousands of samples of African, European, and South Asian ancestry groups from the Trans-Omics for Precision Medicine project and the United Kingdom Biobank. Among the significant loci, many signals of recent selection are shared across ancestry groups. One shared selection signal at a skeletal cell development gene is extremely strong in African ancestry samples.

**Highlights:**

- We propose a method to address multiple testing when scanning along the genome for excess identity-by-descent rates.
- In whole genome simulations, we calculate that the family-wise error rates of our method are close to the desired family-wise significance level.
- We perform six selection scans in two consortium datasets covering different ancestry groups and reference genome builds.
- For a genomic region on chromosome 16, we report extremely high identity-by-descent rates in African ancestry groups and replication in European and South Asian ancestry groups.

## 1. Introduction

Positive natural selection is suggested to be the primary mechanism of phenotypic adaptation [1]. Many reported instances of positive selection in human populations concern adaptive evolution on immunity-related genes [2, 3]. There is also evidence in bacterial, parasite, and insect vector populations for genic selection to evade public health efforts [4, 5, 6]. These examples indicate that the adversarial dynamics between macro-organisms and their microbial pathogens may be a powerful force driving genetic changes in populations. Learning about these genetic changes could be helpful in the design of new vaccines, therapeutics, and interventions in the environment.

Decades of genetics and evolution research have provided many methods to detect positive selection. In general, a statistic is devised to capture different alternative hypotheses from the neutral theory of Kimura [7] or the slightly deleterious theory of Ohta [8], and then the statistic is calculated across the genome to scan for significant evidence against a null model. Some examples of alternative models are selective sweeps [5, 9, 10, 11, 12] and balancing selection [13]. Vitti et al. [1] and Temple et al. [14] categorize these methods into several groups: amino acid substitution rates [15, 16], population differentiation [17, 18], frequency [19, 20, 21], linkage disequilibrium (LD) [22, 23, 24, 25, 26, 27, 28, 29, 30, 14, 31], coalescent [32, 33, 34, 35], approximate Bayesian computation [36], time series [37, 38], and machine learning-based methods [39, 40, 41, 42, 43, 44]. On the one hand, these methods are designed to detect natural selection at different evolutionary timescales or under different mechanisms. On the other hand, the lack of statistical models may have led to the development of many *ad hoc* summary statistics [45]. For instance, some methods clarify that summary statistics a few standard deviations above a genome-wide mean do not have p values [22, 31], and equally so, no adjustment for multiple testing.

We aim to develop a hypothesis testing framework for the selection statistic proposed in Browning and Browning [24] and studied in Temple et al. [14]. One major approach to developing multiple-testing adjustments is to control the family-wise error rate (FWER). FWER is the probability of rejecting the null hypothesis one or more times when the null hypothesis is true [46], which is more conservative than other approaches such as control of false discovery rate [47]. The fundamental question of this article is the following: does our multiple-testing adjustment control the FWER? We give the opinion that rejecting a null hypothesis of neutral evolution and possibly supposing an alternative hypothesis of adaptive evolution is a strong conclusion that warrants conservatism. Hence, we will derive FWER-based multiple-testing corrections.

The p value threshold of 5e-8 is commonly used in genome-wide association studies (GWAS). Based on an assessment of the number of effective hypothesis tests in human genotype array data from the early 2000s, the 5e-8 genome-wide significance level comes from the Bonferroni correction at the 0.05 significance level [48]. Some population genetics studies use this *de facto* significance level even though their study designs and data are different from the human genetics studies in the early 2000s. For instance, in their selection tests, Field et al. [20] and Speidel et al. [33] use the 5e-8 p value threshold.

Permutation or simulation-based approaches can provide interpretable p values and control the FWER under valid permutation or simulation frameworks, but these procedures can be computationally intensive and challenging to design [49, 50, 51, 52, 53, 54]. To remain feasible, some of these simulation-based approaches were applied to sample sizes less than a few thousand [49, 53], or they leveraged the fact that Wald and score statistics from linear models are asymptotically normally distributed [50, 52]. Implementing a simulation-based approach can be infeasible for selection tests that are already computationally intensive in one scan.

Another approach is to model the test statistics under the null hypothesis as a stochastic process and use the properties of that process to determine the threshold. In an identity-by-descent (IBD) mapping study, Browning and Thompson [49] approximate transitions between IBD and non-IBD states as a Markov process and derive an analytical genome-wide significance threshold under their model. In an admixture mapping study, Grinde et al. [52] approximate their Wald test statistics as an Ornstein-Uhlenbeck (OU) process and then calculate the genome-wide significance level with an analytical solution [55, 54]. The Siegmund and Yakir [54] calculation of the genome-wide significance level applies to any scan that can be reasonably modeled as an OU process.

Multiple testing addresses scientific discovery in a single study, whereas much of the consensus scientific progress comes from replicated findings. For example, most scans for recent positive selection in European ancestry populations detect the *LCT* signal [56], which can be as large as thirty-five standard deviations greater than the median of a genome-wide scanning statistic [14]. Indeed, many scans have detected several overlapping selection signals in European ancestry populations [24, 38, 37, 28, 32, 33, 31]. Fewer studies have explored recent positive selection in non-European ancestry populations. Albrechtsen et al. [23] identify the major histocompatibility complex (*MHC*) region as having extreme rates of alleles inferred to be IBD in all human populations. Taliun et al. [57] use the Field et al. [20] method to identify a few loci putatively under recent selection in African and East Asian ancestry samples. In yet another example, Granka et al. [58] enumerate some extreme values of the cross-population extended haplotype homozygosity statistic [17] found in African ancestry populations, but without a multiple-testing adjustment, they exercise caution in the interpretation of their findings. Temple et al. [14] advise that analyzing selection in non-European ancestry samples should proceed with multiple-testing adjustments.

To control the FWER when scanning the genome for excess IBD rates, we propose analytical and simulation-based significance thresholds from an estimated OU process model [24, 14]. We show that the adjusted significance thresholds should approximately control the FWER under some central limit theorem conditions [59]. The IBD rate scan is computationally efficient; hence, we can measure its FWER in simulation studies. We also demonstrate the effects of various analysis decisions on the empirical FWER and statistical power, including user-defined centiMorgan (cM) spacings and IBD segment detection thresholds. We show that the heuristic four standard deviations above the autosome-wide median threshold used in the Browning and Browning [24] and Temple et al. [14] studies may have been reasonable for European ancestry populations but that the genome-wide significance threshold should be more stringent for some African populations. We detect a statistically significant locus in two African ancestry sample sets whose excess IBD rates are more than ten standard deviations above the respective genome-wide means and which replicates in European and South Asian ancestry samples. Nevertheless, after adjusting for multiple testing, we observe less than twelve signals of recent positive selection in any given cohort.

## 2. Materials and Methods

### 2.1. Hypothesis testing framework

First, we define the implicit hypothesis test in the IBD rate scan [24, 14]. When modeling the spatial process, we use the same mathematical notation as Temple and Thompson [59] with minor revisions. Let *Y_a,b_*(*m*) be the indicator that the IBD segment between haplotypes *a* and *b* is longer than a detection threshold and overlaps the *m*^th^ focal position. The IBD rate at the *m*^th^ locus is Ў *_m_* = *f* (*n*)^−1^ Σ_(_*_a,b_*_)_ *Y_a,b_*(*m*), where *f* (*n*) = 2*n*(2*n* − 1)*/*2 − 2*n* in diploids and *f* (*n*) = tn) in haploids. The hypothesis test we consider is

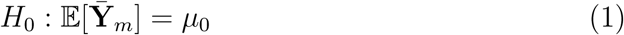

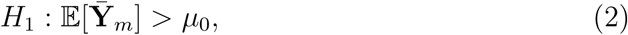

where *µ*_0_ is a genome-wide mean IBD rate around a locus. This null model is consistent with no positive selection. The alternative model is consistent with positive selection *or* other evolutionary mechanisms.

Let *µ*^_1:*M*_ and *σ*^_1:*M*_ be the sample mean and sample standard deviation of *M* IBD rates along the genome:

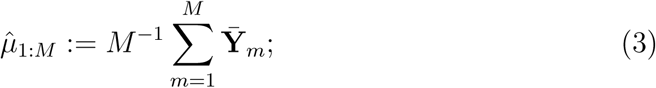

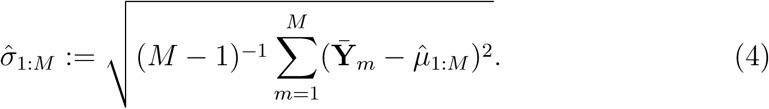

Browning and Browning [24] and Temple et al. [14] suggest a heuristic threshold of *µ*^_1:_*_M_* + 4 × *σ*^_1:_*_M_* as strong evidence against the null model. (They use the genome-wide median, not the mean, which can be more robust to outliers like *LCT* selection.) Under asymptotic conditions on sample size, population demography, and the detection threshold, the standardized IBD rate **Z**^-^*_m_* around the *m*^th^ locus is normally distributed [59]. The heuristic threshold corresponds to a significance level of 1 − Φ(4) = 3.17 × 10^−5^.

We use the same test statistic as Browning and Browning [24] and Temple et al. [14], except we adapt the number of standard deviations to the correlation structure in a distinct sample:

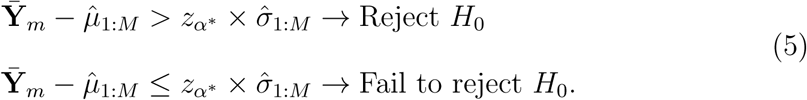

This test corresponds to a one-sample one-sided t test or a z test when the number of tests *M* is large. The significance level *α*^*^ comes from a multiple-testing correction at the family-wise significance level *α*, and *z_α_*\* is the corresponding standard normal quantile.

To determine multiple-testing corrections, we model standardized IBD rates along the genome

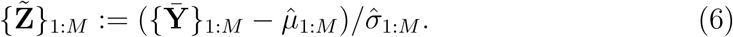

as a correlated OU process. This model has previously been used to determine multiple-testing corrections in admixture mapping [51, 52] and linkage analysis [55]. The OU process is normally distributed at every point, is spatially homogeneous, and has the first-order Markov property. Assuming normality at every point is supported by the Temple and Thompson [59] central limit theorems and may be reasonable in human genetics studies. Spatial homogeneity is an assumption consistent with neutral evolution and uniform IBD segment detection accuracy. Compared to the Grinde et al. [52] admixture mapping statistics, which are provably Markov, the IBD rate along the chromosome is not a Markov process (Temple [60] gives a simple counterexample). Therefore, we assume that the IBD rate process is nearly Markov, at least so much so that the violation does not affect our multiple-testing corrections.

The standard OU process has a specific correlation pattern. Namely, if the genetic distance between consecutive focal positions is set to be constant Δ, then the covariance between standardized IBD rates **Z**^-^*_m_*_1_ and **Z**^-^*_m_*_2_ at different loci is

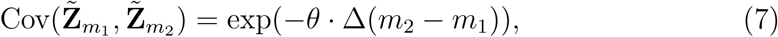

where *θ* is an exponential decay parameter. The exponential decay parameter *θ* is not known for the IBD rate process but must be estimated, whereas *θ* is the time of admixture in Grinde et al. [52], which can be estimated or assumed from prior knowledge.

### 2.2. Multiple-testing corrections

#### 2.2.1. Analytical approximation

To control the FWER, we must determine the multiple-testing quantile *z_α_*\* such that *P* (max*_m_* **Z**^-^*_m_*≥ *z_α_*\*) = *α*. Let *L* be the total length of the genome (in Morgans), *C* be the number of chromosomes, and Φ and *ϕ* be the cumulative distribution and density functions of the standard normal random variable. Siegmund and Yakir [54] provide the FWER-based analytical approximation

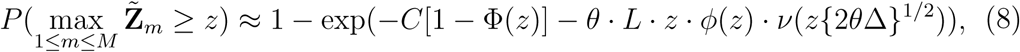

where *ν*(·) accommodates the discretization of the continuous stochastic process. When the Morgan step size Δ → 0 (the continuous process), *ν*(0) = 1. We determine *z_α_*\* from Equation 8 with a root solver, which runs in seconds. This approach is an example of finding the first hitting time of a stochastic process.

#### 2.2.2. Simulation-based approach

Another way to control the FWER is to simulate the OU process for known or estimated *θ*. Let *J* be the number of simulations and *M*:= ⌊*L* ÷ Δ⌋. The simulation approach goes as follows.

Algorithm 1.

1. Let z1:J be an empty vector.
2. For *j* in 1 to *J*:

a. Draw *z*_1_ = *Z*_1_ ∼ *N* (0, 1).
b. For *m* in 2 to *M*:

i. Draw *z_m_* = *Z* | *z_m_*_−1_ ∼ *N* (*z_m_*_−1_ · exp(−*θ* · Δ), 2 − 2 · exp(−*θ* · Δ)).
c. _(c)_ Append max*_m_ z_m_* to the vector **z**_1:_*_J_*
3. Return the (1 − *α*)% quantile of **z**_1:_*_J_*.

For family-wise significance levels like 0.01 or 0.05, this simulation approach requires a few thousand simulations and runs within a few minutes (depending on the genome length *L*). This multiple-testing correction is valid when the true model is the OU process. A precise algorithm would simulate individual OU processes for different chromosome lengths, but for simplicity, we simulate a single chromosome of the total genome length instead.

### 2.3. Estimator of the exponential decay parameter

Before standardizing the IBD rates, we adjust for extreme outliers that could be present in real genetic data. First, we compute an initial genome-wide median IBD rate plus four standard deviations. Second, we compute a revised genome-wide mean IBD rate and standard deviation, excluding the IBD rates that exceed the initial threshold. We standardize the IBD rates with the revised mean and standard deviation estimates. This step is suitable for the reproducible workflow of Temple et al. [14], whereas filtering out known exceptions like *LCT* selection in European ancestry populations is less amenable to automation [61].

To estimate the exponential decay parameter *θ*, we regress estimated autocovariances on genetic position. We apply linear interpolation to the recombination map to hold the spacings between IBD rates constant. Then, we estimate the covariance between standardized IBD rates at genetic positions Δ times some integer constant apart, excluding IBD rates that exceed the initial threshold. The integer scalars increment by one until the covariance is between positions maximum 4.0 cM apart. We fit a simple log-linear model with no intercept, where the integer-scaled Δ’s are the covariates and the estimated autocovariances are the response variables. The fitted slope parameter is an estimator *θ*^^^ of the exponential decay parameter.

### 2.4. Simulating IBD rate processes

#### 2.4.1. Null hypothesis model

We evaluate control of the FWER and the accuracy of our estimator *θ*^^^ with large-scale coalescent simulations. We use msprime [62] to simulate ten chromosomes, each of length 100 cM, and we use tskibd [6] to get IBD segment lengths longer than 2.0 and 3.0 cM from the tree sequence output by msprime. The constant recombination rate is 1e-8. We consider previously defined demographic scenarios of a population bottleneck, a constant population of size fifty thousand individuals, and staged exponential growth [60, 59, 63, 14]. The demographic scenario affects the exponential decay parameter *θ*. Unless otherwise specified, our default demographic scenario is the population bottleneck.

We estimate *θ* from the autocovariances of simulated IBD segments, and then we use the estimate *θ*^^^ to calculate our multiple-testing adjusted thresholds. For these calculations of the genome-wide significance level, we consider different step sizes 0.02, 0.05, and 0.10 cM. Unless otherwise specified, the default step size is 0.02 cM. The estimator *θ*^^^ should be agnostic to the cM spacing, but the genome-wide significance level should decrease monotonically with the cM spacing.

To empirically measure the FWER, we consider five hundred simulations of entire genomes from twenty-five hundred diploids. FWER is calculated as the percentage of the five hundred null model simulations with at least one significant result. We explore the family-wise significance levels of 0.01, 0.05, and 0.10. Unless otherwise specified, we use the 0.05 family-wise significance level. We use the discrete-spacing analytical approximation as our default multiple-testing correction.

The data for our simulations amounts to 1 terabyte (TB) compressed disk storage, predominantly due to the msprime tree sequences. We are unable to make VCF marker data for all our simulations and, therefore, to infer IBD segments, which would create many more TB of additional disk memory. In Appendix A.1, we analyze the accuracy of IBD segment detection in VCF marker data.

#### 2.4.2. Selective sweep alternative model

To calculate statistical power, we consider hard sweeps as the alternative model. This evolutionary scenario concerns a single advantageous allele increasing in frequency, with the rate of change parameterized by the selection coefficient *s* [64, 65, 66]. For the population bottleneck and staged exponential growth scenarios, we simulate IBD segments overlapping a focal point for hard sweeps with *s* ≥ 0.006 and current-day allele frequency *p*(0) = 0.25, 0.50, 0.75 with the Temple et al. [63] algorithm. Based on the results of Temple et al. [14], we believe that the algorithm in Temple et al. [63] simulates IBD rates around a locus similar to those drawn from tree sequences by tskibd, which itself has not been independently benchmarked. For the constant population size scenario, we do consider tree sequences, and therefore tskibd segments, simulated with positive selection, which is an msprime feature only available for constant populations [62, 67].

Power is calculated as the proportion of our selective sweep simulations (alternative hypotheses) in which we reject the null model. The threshold in our power calculations is the average of the multiple-testing adjusted thresholds in our five hundred neutral simulations. We estimate power using two hundred simulations for each pair of selection coefficient and current-day sweeping allele frequency.

### 2.5. Pre-processing genetic data

In our study, we focus on selection scans in African, European, and South Asian ancestry groups from the Trans-Omic for Precision Medicine (TOPMed) project [57] and the United Kingdom Biobank (UKBB) [68]. The TOPMed data that we analyze includes more than thirty thousand whole genome sequences from multiple ethnic groups represented in the United States of America, combining samples from various cohort studies. We use the 318,858,817 filtered autosomal markers from the TOPMed data phased with Beagle 5.2 in Browning et al. [69]. UKBB is a biomedical database containing genotype array data from nearly five hundred thousand participants between 40 and 69 years of age. We use the 711,651 filtered autosomal markers from the UKBB SNP array data in Browning et al. [69]. The TOPMed and UKBB datasets are kept separate in all analyses.

#### 2.5.1. Trans-Omics for Precision Medicine

We analyze the whole genome sequences of multiple ancestry groups inferred by Temple et al. [14]. These ancestry groups were defined by principal component analysis (PCA) [70, 71] and validated with ADMIXTURE [72]. Individuals inferred to be third-degree or closer relatives are excluded [14]. One of our subsets is the 13,778 European ancestry samples studied by Temple et al. [14], which we now refer to as the EUR1 ancestry group.

Another European ancestry group we define is EUR2, comprising 1719 samples whose principal components are near to but distinct from those of the samples in the EUR1 group. Sixty-four percent of these samples come from the BioMe Biobank cohort study at Mt. Sinai School of Medicine in New York City, which is a dataset known to contain many samples inferred to have Ashkenazi Jewish ancestry [73]. For this group, we infer a demographic history that sharply drops to an effective size as small as one thousand in the most recent thirty generations (IBDNe using ≥ 2.0 cM IBD segments [74]). In an Ashkenazi Jewish sample,

Carmi et al. [75] infer a recent bottleneck of the effective size of a few hundred diploids, which Tian et al. [76] say is consistent with their demographic inference of a Framingham Heart Study subset. Carmi et al. [75] state that the Ashkenazi Jewish population is most genetically similar to European and Middle Eastern populations, which is consistent with the Temple et al. [14] principal components analysis and the fastSTRUCTURE analysis [77] done by Wu et al. [73].

Using the first principal component, we define an inferred African ancestry group (AFR) of 1737 samples. Based on the ADMIXTURE validation study of Temple et al. [14], these samples have minimum and mean global ancestry proportions of 0.88 and 0.93 with respect to the Yoruba in Ibadan, Nigeria (YRI) reference panel [78, 79]. Fifty-four percent of these samples self-report as Black or African American, and forty-six percent self-report as Other. Only samples from the Barbados Asthma Genetics Study (BAGS), Jackson Heart Study (JHS), and Hypertension Genetic Epidemiology Network Study (HyperGen) cohorts are represented in this subset. Afro-Caribbeans living in Barbados are in the BAGS study, whereas African Americans living in the southern continental United States are in the JHS and HyperGen studies.

To detect IBD segments in the TOPMed sample sets, we use the algorithm parameters in the Temple et al. [14] workflow. In the EUR1 ancestry group, we use the IBD segments previously inferred by Temple et al. [14]. We perform preliminary analyses of chromosomes 19 to 22 with ibd-ends [24] to get estimates of the error rate parameter, eventually specifying the error rate err=1.5e-4 for all three groups. All TOPMed analyses use the 2019 pedigree-based genetic map from deCODE Genetics [80]. This recombination map is aligned to the GRCh38 reference genome.

#### 2.5.2. United Kingdom Biobank

We also analyze subsets of the UKBB samples who self-report as various non-white ethnic groups. The first subset includes 5660 individuals who self-report as Indian British [68]. The second subset consists of 3202 individuals who self-report as Black British (African in Bycroft et al. [68]). We phased the sample sets individually with Beagle version 5.4. Based on genetic relatedness inference in Cai et al. [81], we remove closely related individuals from both subsets, resulting in 5374 Indian British and 3146 Black British samples.

We also analyze the 408,891 UKBB white British samples previously studied in Browning and Browning [24]. (The group definition ‘white’ comes from a combination of self-reported British ethnic background and similar scores in a PCA analysis [68].) The SNP array data was previously phased with Beagle 5.2 as described in Browning et al. [69].

To detect IBD segments in the UKBB sample sets, we modify our hap-ibd settings to min-seed=1.8, min-extend=0.5, min-output=1.8, and a minor allele frequency of 0.001. We have not explored the accuracy of these settings in simulated array data. Still, we show in our results that our analyses of array data are consistent with our analyses of sequence data and with the existing literature on some selected loci. In the white British, Indian British, and Black British groups, we perform preliminary analyses of chromosomes 19 to 22 with ibd-ends to get estimates of the error rate parameter, eventually specifying the error rate err=3.0e-4 for all groups. All UKBB analyses use the Bhérer et al. [82] pedigree-based genetic map. This recombination map is aligned to the GRCh37 reference genome.

## 3. Results

### 3.1. Simulated Ornstein-Uhlenbeck processes

We conduct a simple validation study to determine if the discrete-spacing analytical approximation and simulation-based genome-wide significance levels control the FWER when data is simulated from an OU process. The simulation settings are in Figures S1 and S2. Figure S1 shows that estimates *θ*^^^ of the exponential decay parameter are approximately equal to the true value when 30 ≤ *θ* ≤ 90 and the genome size is ≥ 400 cM. Figure S2 shows FWERs using the multiple-testing corrections at a family-wise significance level of 0.05. The FWERs from the discrete-spacing analytical approach are between 0.04 and 0.05 and less than 0.03 when *θ* ≥ 30 and *θ* = 1, respectively. Grinde et al. [52] also find that the discrete-spacing analytical approximation is conservative when *θ* ≈ 10. The FWERs from the simulation-based approach are approximately 0.05 for all *θ*. We thus recommend using the simulation-based approach if *θ* ≤ 20. While the discrete-spacing analytical approach may be slightly conservative compared to the simulation-based approach, simulating 500 OU processes of size equal to the 22 human autosomes can take as much as ten minutes.

### 3.2. Simulated IBD rate processes

#### 3.2.1. Estimating the exponential decay parameter

Box plots in Figure S3 show the percentiles of estimates *θ*^^^ using IBD segments ≥ 2.0 and ≥ 3.0 cM from data simulated under the null hypothesis with the population bottleneck scenario. Regardless of the step size Δ, the distribution of estimates *θ*^^^ is the same, which is expected. The medians of estimates *θ*^^^ for the ≥ 2.0 and ≥ 3.0 cM processes are roughly 40 and 62.5, respectively. As *θ* increases, and holding the genetic distance between two positions constant, the covariance between the two IBD rates decreases, which could be interpreted as fewer detectable IBD segments overlapping nearby loci on average. Estimates for *θ* are smaller in the ≥ 2.0 cM scan versus the ≥ 3.0 cM scan because an IBD segment ≥ 2.0 cM is less likely to also overlap the next focal point than an IBD segment ≥ 3.0 cM is.

For the staged exponential growth scenario, the medians of estimates *θ*^^^ are 74.75 and 56.78 for the ≥ 2.0 and ≥ 3.0 cM IBD rate processes, respectively. For the population of constant size fifty thousand diploid individuals, the medians of estimates *θ*^^^ are 58.84 and 44.97 for the ≥ 2.0 and ≥ 3.0 cM IBD rate processes, respectively. We expect different true *θ* and therefore different estimates *θ*^^^ because demography influences the IBD segment length distribution [74, 81, 60].

#### 3.2.2. Family-wise error rates

Table 1 reports the multiple-testing adjusted significance levels and the empirical FWERs for the discrete-spacing analytical approximation and simulation-based approaches in the ≥ 2.0 cM IBD rate processes. The adjusted significance levels from the analytical and simulation-based approaches are nearly an order of magnitude larger than those using the Bonferroni correction. At the 0.05 family-wise significance level, the FWERs of our analytical and simulation-based approaches are inflated by more than 150%. In contrast, the FWERs of the Bonferroni method with testing every 0.02 cM are deflated by less than 50%. At the 0.10 family-wise significance level, the average standard deviations above the mean of the analytical and simulation-based approaches are 4.196 and 4.176. Temple and Thompson [59] give one plausible explanation for the anti-conservativeness of the hypothesis test with ≥ 2.0 cM segments, which is that the upper tail of the IBD rate’s distribution may be heavier than the upper tail of a normal distribution.

**Table 1:** Genome-wide significance levels and family-wise error rates after multiple-testing corrections. Family-wise significance levels are adjusted for multiple testing based on scans over 10 chromosomes of size 100 cM and tests every 0.02 cM (50,000 total tests). The multiple-testing analytical and simulation-based thresholds are based on a fitted Ornstein-Uhlenbeck process. Family-wise error rate (FWER) is the percentage of five hundred genome-wide scans with at least one statistically significant result. The demographic scenario is the population bottleneck. The IBD segment detection threshold is 2.0 cM.

| **Family-wise**<br>**level** | **Genome-wide**<br>**Analytical** | **Simulation** | **Bonferroni** | **FWER**<br>**Analytical** | **Simulation** |
| --- | --- | --- | --- | --- | --- |
| 0.01 | 1.08e-6 | 1.30e-6 | 2.08e-7 | 0.024 | 0.028 |
| 0.05 | 6.24e-6 | 7.03e-6 | 1.04e-6 | 0.088 | 0.098 |
| 0.10 | 1.36e-5 | 1.49e-5 | 2.08e-6 | 0.140 | 0.146 |

Table S1 reports the adjusted significance levels and FWERs of the multiple-testing approaches using the 3.0 cM threshold. In this case, the IBD rate overlapping a locus may be better approximated by a normal distribution than in the ≥ 2.0 cM selection scan (conditions on the detection threshold in Temple and Thompson [59]). The FWERs of the analytical and simulation-based approaches are indeed conservative in the ≥ 3.0 cM excess IBD rate scan. We thus remark that there are two counteracting factors affecting FWER control: the multiple-testing adjustments are conservative in true OU processes (Figure S2), but the test could be anti-conservative if the OU process is a poor approximation for the IBD rate process.

For the anti-conservative ≥ 2.0 cM excess IBD rate scan, we consider modifying the test to explore whether the significant results *barely* exceed the threshold. We calculate at each locus the minimum of its value and the flanking values to its left and right. Next, we calculate the maximum over the entire genome of these aggregated minimum values:

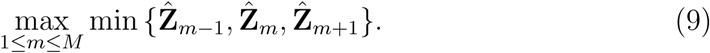

Figure 1 shows that FWERs decrease when using the max-min statistic. This result indicates that a considerable proportion of the family-wise errors correspond to marginally significant results.

**Figure 1:**
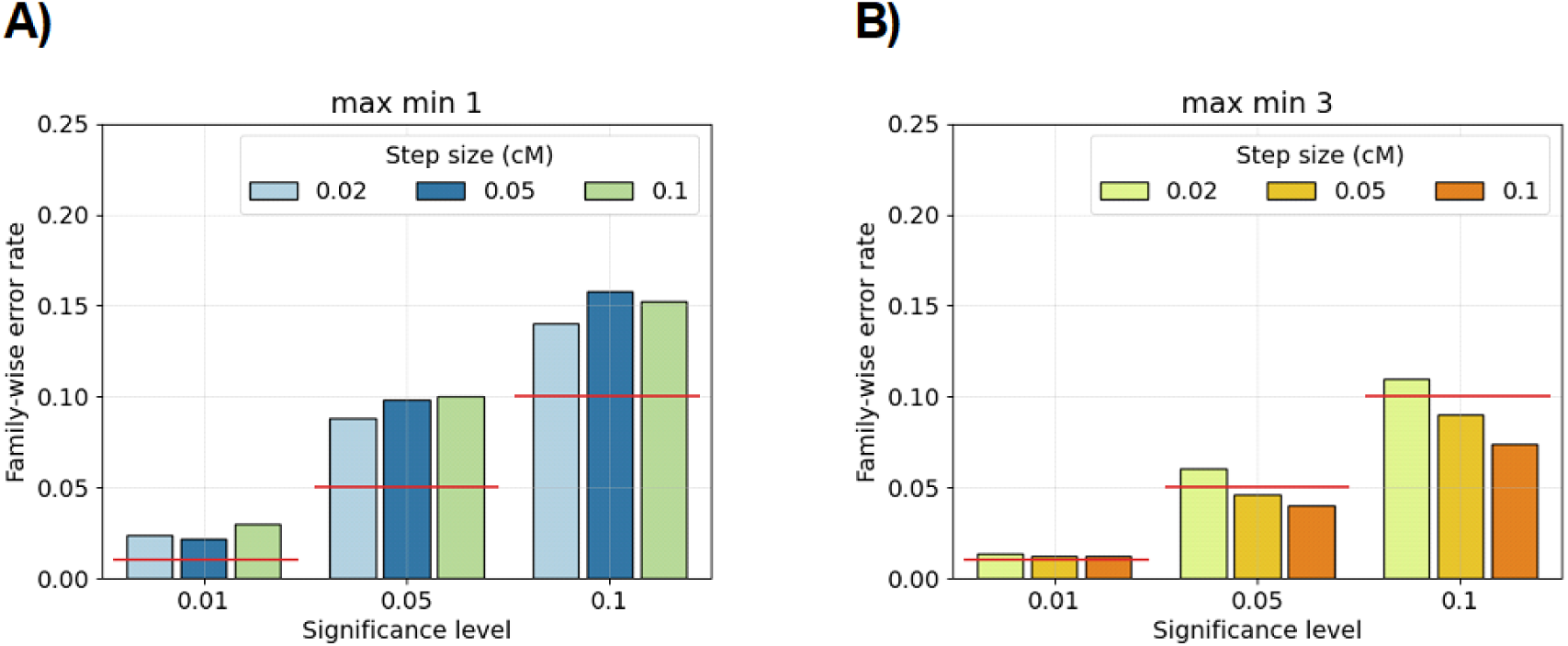
Family-wise error rates for genome-wide hypothesis testing in null model simulations. Bar plots show family-wise error rates (y-axis) using true IBD segments ≥ 2.0 cM from simulated IBD rate processes. The hypothesis testing method is the discrete-spacing analytical approximation. In each non-overlapping window of size A) 1 or B) 3 marginal test statistics, we compute the minimum of IBD rates at each step, and the test is if the maximum over all windows is less than or greater than the multiple-testing quantile. There are five hundred simulations for each combination of significance level (x-axis) and step size (colors in legend). Family-wise significance levels are denoted with horizontal red lines. The demographic model is the population bottleneck. The amount of data for each simulation is equal to ten chromosomes of uniform length 100 cM.

Next, when there is a significant result, we investigate how many significant results there are. Since the IBD rate process has non-negligible correlations, we anticipate multiple significant results adjacent to each other. Across non-overlapping windows of varying sizes, we count the number of windows that have a significant result. Figure S4 shows that the number of windows with a significant result decreases to a median of 1 when the window size is ≥ 0.20 cM and the family-wise significance level is ≤ 0.05. Altogether, we tend to find only one or a few marginally significant results in aggregated regions less than 0.5 cM when a Type 1 error is made.

For the staged exponential growth scenario, the average standard deviations above the mean using the discrete-spacing analytical approximation are 4.00 and 4.35 for the ≥ 2.0 and ≥ 3.0 cM IBD rate processes, respectively. The average genome-wide significance levels are 5.41e-6 and 6.82e-6, and the FWERs are 0.148 and 0.036. For the population of constant size fifty thousand diploid individuals, the average quantiles using the analytical approximation are 4.36 and 4.31 for the ≥ 2.0 and ≥ 3.0 cM IBD rate processes, respectively. The average genome-wide significant levels are 6.56e-6 and 8.38e-6, and the FWERs are 0.114 and 0.034. Regardless of the demographic scenario, the ≥ 2.0 and ≥ 3.0 cM scans may have anti-conservative and conservative control of the FWER, respectively.

#### 3.2.3. Statistical power in selective sweeps

Figures 2 and S6A show the power estimates for the ≥ 2.0 cM IBD rate scan in the population bottleneck, staged exponential growth, and constant population size scenarios with selection coefficients 0.006 ≤ *s* ≤ 0.014 and current-day allele frequencies 0.25 ≤ *p*(0) ≤ 0.75. Power estimates are uniformly greater with the current-day allele frequency *p*(0) = 0.50 as opposed to *p*(0) = 0.25 or *p*(0) = 0.75. The increased ability to detect positive selection when the sweep is at an intermediate present-day frequency is consistent with the analyses in Temple et al. [14]. For the population bottleneck simulations, power estimates are less than 5% when *s* ≤ 0.008 but greater than 90% when *s* ≥ 0.014. In between these extremes, power estimates range from 15% to 40% when *s* = 0.010 and from 55% to 85% when *s* = 0.012. Depending on *s* and *p*(0), power estimates are 10% to 30% higher in the staged exponential growth simulations than they are in the population bottleneck simulations. In constant population size simulations, power estimates are between 0% and 10% when *s* ≤ 0.012 but as high as 40% when *s* = 0.014 and *p*(0) = 0.50. The parameter boundary *s* ≤ 0.01 and *s >* 0.01 marks a transition consistent across all our demographic scenarios when the ≥ 2.0 cM scan has some nonzero statistical power.

**Figure 2:**
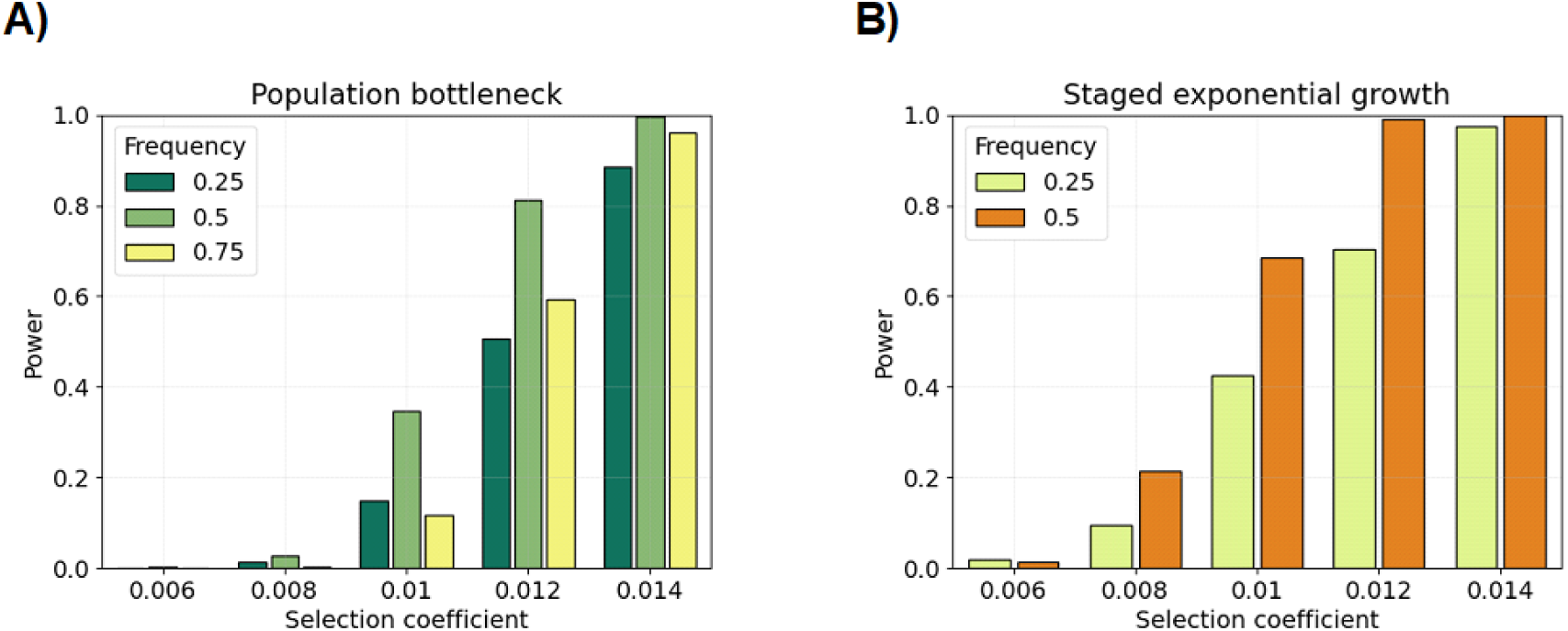
Power simulations for different selection coefficients and current-day sweeping allele frequencies. Bar plots show statistical power (y-axis) using true IBD segments ≥ 2.0 cM overlapping the selected allele in the A) population bottleneck and B) staged exponential growth demographic scenarios. Hypothesis testing is based on the discrete-spacing analytical threshold with the step size 0.02 cM. Power is the proportion of tests where the null model is rejected at the p value threshold corresponding to the 0.05 family-wise significance level. There are two hundred simulations for each pair of selection coefficient (x-axis) and current-day allele frequency (colors in legend).

Figures S5 and S6B show the power estimates for the ≥ 3.0 cM scan in the population bottleneck, staged exponential growth, and constant population size scenarios. In the population bottleneck and constant population size simulations, we measure zero power for all combinations of selection coefficients and allele frequencies. In the staged exponential growth simulations, we measure power between 10% and 50% for selection coefficients *s >* 0.01 and zero for selection coefficients *s* ≤ 0.008. Regardless of demography, rejecting the null model in the ≥ 3.0 cM scan could be evidence of an exceptionally strong sweep.

### 3.3. Multiple-testing corrections for human ancestry groups

We modify the Temple et al. [14] workflow to incorporate the analytical approximation and simulation-based approaches for multiple testing. We also provide genome-wide significance levels under the null model that IBD rates are normally distributed. (IBD rates are asymptotically normally distributed under some conditions on large sample size and population size [59].)

For each sample set, we compute IBD rates every 0.02 cM for IBD segments ≥ 2.0 and ≥ 3.0 cM. Figure S7 indicates that the empirical distributions of IBD rates around a locus resemble normal distributions in our sample sets. Figure S8 shows the estimated autocovariances and fitted exponential curve for all our ancestry and ethnicity groups. Upon visual inspection, the fitted exponential curves match the chromsome-specific autocovariances well in the plots for the European ancestry and UKBB Indian British sample sets. In the TOPMed AFR ancestry and UKBB Black British groups, the fitted exponential curves fit the long-range autocovariances well but not the short-range autocovariances.

For IBD segments ≥ 2.0 cM, the exponential decay parameter estimates *θ*^^^ are 45, 30, 50, 49, 83, and 78 for the TOPMed EUR1 ancestry, TOPMed EUR2 ancestry, UKBB white British 410k, UKBB Indian British, TOPMed AFR ancestry, and UKBB Black British groups, respectively. The corresponding discrete-spacing analytical thresholds are IBD rates 1.94e-4, 5.89e-3, 2.66e-4, 1.82e-4, 2.64e-4, and 3.55e-4, and the corresponding genome-wide significance levels are 2.27e-6, 3.27e-6, 2.13e-6, 2.16e-6, 1.36e-6, and 1.46e-6. For each of these estimates of the exponential decay parameter, the discrete-spacing analytical and simulation-based approaches should provide similar genome-wide significance levels (Figure S2).

For IBD segments ≥ 3.0 cM, the exponential decay parameter estimates *θ*^^^ are 33, 36, 39, 53, and 45 for the TOPMed EUR1 ancestry, UKBB white British 410k, UKBB Indian British, TOPMed AFR ancestry, and UKBB Black British groups, respectively. The corresponding discrete-spacing analytical thresholds are IBD rates 4.49e-5, 8.08e-5, 9.31e-5, 6.10e-5, and 8.07e-5, and the corresponding genome-wide significance levels are 3.05e-6, 2.87e-6, 2.70e-6, 2.02e-6, and 2.35e-6.

### 3.4. Selection scans for human ancestry groups

Figure 3 shows the ≥ 2.0 cM IBD rates along the autosomes, the autosome-wide median, the heuristic four standard deviations above the median threshold, and the multiple-testing adjusted thresholds for the TOPMed EUR1 ancestry, UKBB white British, UKBB Indian British, and TOPMed EUR2 ancestry groups. Figure 4 shows the ≥ 2.0 cM IBD rates along the autosomes, the autosome-wide median, the heuristic four standard deviations above the median threshold, and the multiple-testing adjusted thresholds for the TOPMed AFR ancestry and UKBB Black British groups. In Tables 2 and 3, we report loci where IBD rates exceed the genome-wide significance threshold for a contiguous stretch of 0.50 cM. We annotate loci with genes or gene complexes if they have been previously reported in the literature, are shared across analyses, or contain only a couple of genes. We calculate p values under the null model for the position in a region with the highest IBD rate.

**Figure 3:**
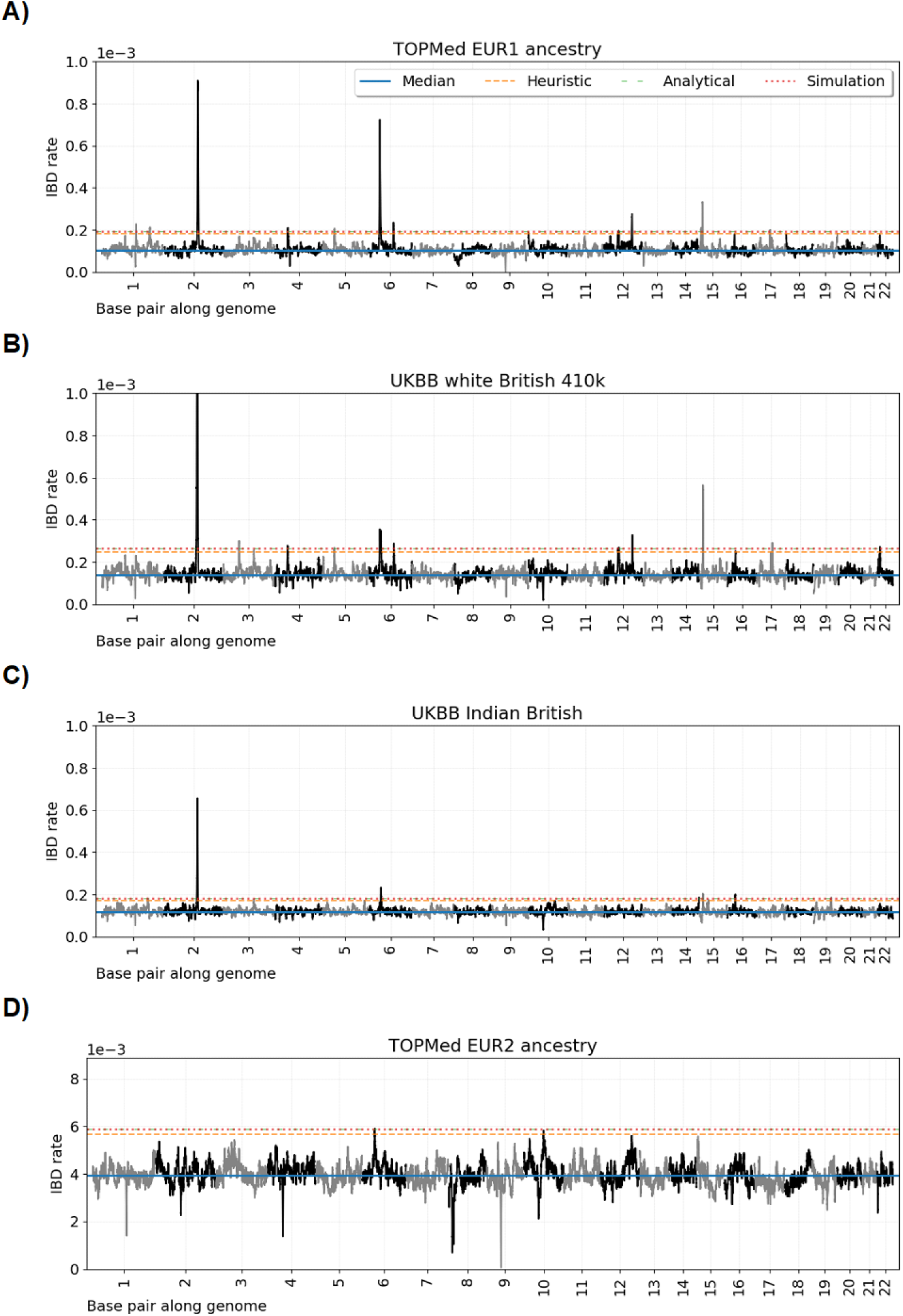
Genome-wide IBD rate scans in European ancestry and Indian British samples. Line plots show IBD rates (y-axis) every 0.02 cM along the twenty-two human autosomes. The dataset analyzed is given in the subplot titles. Horizontal lines show (blue) the genome-wide median IBD rate, (orange) the heuristic threshold of four standard deviations above the median IBD rate, (green) the discrete-spacing analytical threshold, and (red) the simulation-based threshold. The analytical and simulation-based thresholds are less than 5e-6 apart.

**Figure 4:**
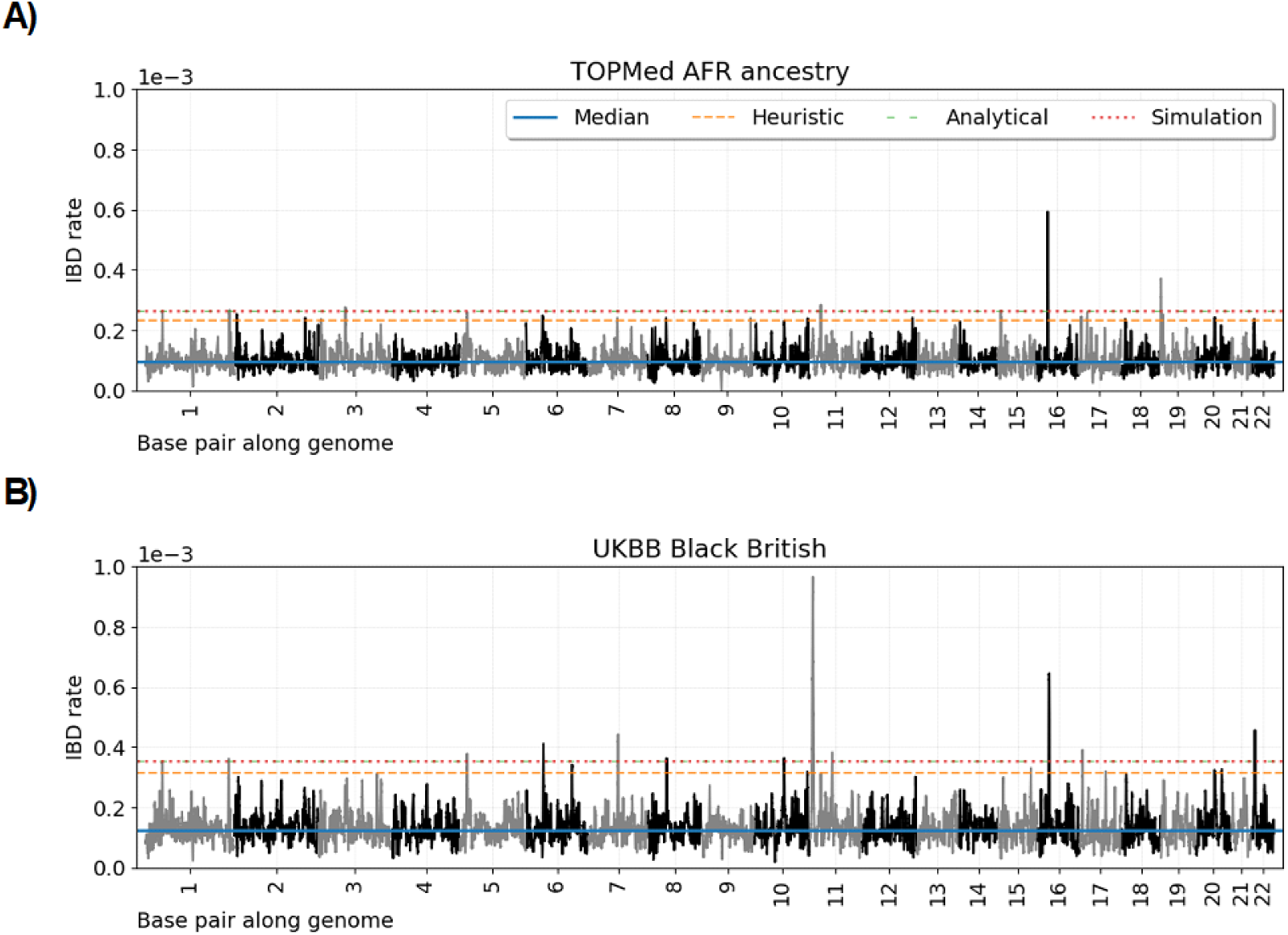
Genome-wide IBD rate scans in African ancestry and Black British samples. Line plots show IBD rates (y-axis) every 0.02 cM along the twenty-two human autosomes. The dataset analyzed is given in the subplot titles. Horizontal lines show (blue) the genome-wide median IBD rate, (orange) the heuristic threshold of four standard deviations above the median IBD rate, (green) the discrete-spacing analytical threshold, and (red) the simulation-based threshold. The analytical and simulation-based thresholds are less than 5e-6 apart.

**Table 2:**
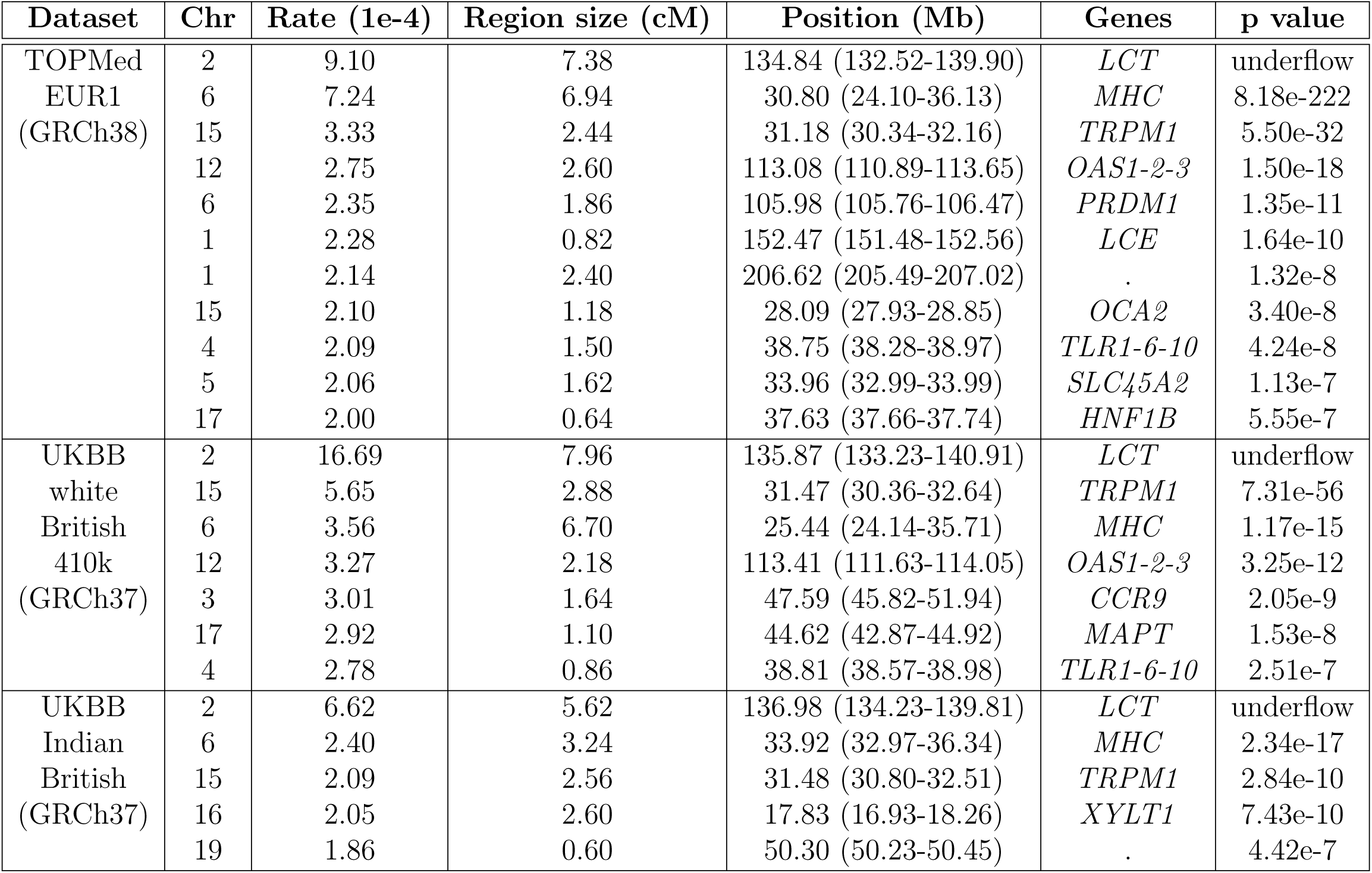
Loci detected in European ancestry and UKBB Indian British selection scans. We report loci where identity-by-descent (IBD) rates exceed the multiple-testing analytical thresholds of 1.94e-4, 2.66e-4, and 1.82e-4 for the TOPMed EUR1 ancestry, UKBB white British, and UKBB Indian British sample sets, respectively. The maximum IBD rate is given for each locus. Physical positions for the location of the maximum IBD rate and the span of excess IBD rates are shown in megabases (Mb). We report the size in centiMorgan (cM) of each region, which is defined to be a contiguous stretch of IBD rates exceeding the genome-wide significance threshold. Pedigree-based recombination maps from Halldorsson et al. [80] and Bhérer et al. [82] aligned to the GRCh38 and GRCh37 reference genomes are used for inferring IBD segments in the TOPMed and UKBB sample sets, respectively. p values are calculated assuming the null model that IBD rates are normally distributed. Annotated genes or gene complexes are discussed in the main text. The IBD segment detection threshold is 2.0 cM.

| Dataset | Chr | Rate (1e-4) | Region size (cM) | Position (Mb) | Genes | p value |
| --- | --- | --- | --- | --- | --- | --- |
| TOPMed<br>EUR1<br>(GRCh38) | 2 | 9.10 | 7.38 | 134.84 (132.52-139.90) | *LCT* | underflow |
|  | 6 | 7.24 | 6.94 | 30.80 (24.10-36.13) | *MHC* | 8.18e-222 |
|  | 15 | 3.33 | 2.44 | 31.18 (30.34-32.16) | *TRPM1* | 5.50e-32 |
|  | 12 | 2.75 | 2.60 | 113.08 (110.89-113.65) | *OAS1-2-3* | 1.50e-18 |
|  | 6 | 2.35 | 1.86 | 105.98 (105.76-106.47) | *PRDM1* | 1.35e-11 |
|  | 1 | 2.28 | 0.82 | 152.47 (151.48-152.56) | *LCE* | 1.64e-10 |
|  | 1 | 2.14 | 2.40 | 206.62 (205.49-207.02) | . | 1.32e-8 |
|  | 15 | 2.10 | 1.18 | 28.09 (27.93-28.85) | *OCA2* | 3.40e-8 |
|  | 4 | 2.09 | 1.50 | 38.75 (38.28-38.97) | *TLR1-6-10* | 4.24e-8 |
|  | 5 | 2.06 | 1.62 | 33.96 (32.99-33.99) | *SLC45A2* | 1.13e-7 |
|  | 17 | 2.00 | 0.64 | 37.63 (37.66-37.74) | *HNF1B* | 5.55e-7 |
| UKBB<br>white<br>British<br>410k<br>(GRCh37) | 2 | 16.69 | 7.96 | 135.87 (133.23-140.91) | *LCT* | underflow |
|  | 15 | 5.65 | 2.88 | 31.47 (30.36-32.64) | *TRPM1* | 7.31e-56 |
|  | 6 | 3.56 | 6.70 | 25.44 (24.14-35.71) | *MHC* | 1.17e-15 |
|  | 12 | 3.27 | 2.18 | 113.41 (111.63-114.05) | *OAS1-2-3* | 3.25e-12 |
|  | 3 | 3.01 | 1.64 | 47.59 (45.82-51.94) | *CCR9* | 2.05e-9 |
|  | 17 | 2.92 | 1.10 | 44.62 (42.87-44.92) | *MAPT* | 1.53e-8 |
|  | 4 | 2.78 | 0.86 | 38.81 (38.57-38.98) | *TLR1-6-10* | 2.51e-7 |
| UKBB<br>Indian<br>British<br>(GRCh37) | 2 | 6.62 | 5.62 | 136.98 (134.23-139.81) | *LCT* | underflow |
|  | 6 | 2.40 | 3.24 | 33.92 (32.97-36.34) | *MHC* | 2.34e-17 |
|  | 15 | 2.09 | 2.56 | 31.48 (30.80-32.51) | *TRPM1* | 2.84e-10 |
|  | 16 | 2.05 | 2.60 | 17.83 (16.93-18.26) | *XYLT1* | 7.43e-10 |
|  | 19 | 1.86 | 0.60 | 50.30 (50.23-50.45) | . | 4.42e-7 |

**Table 3:**
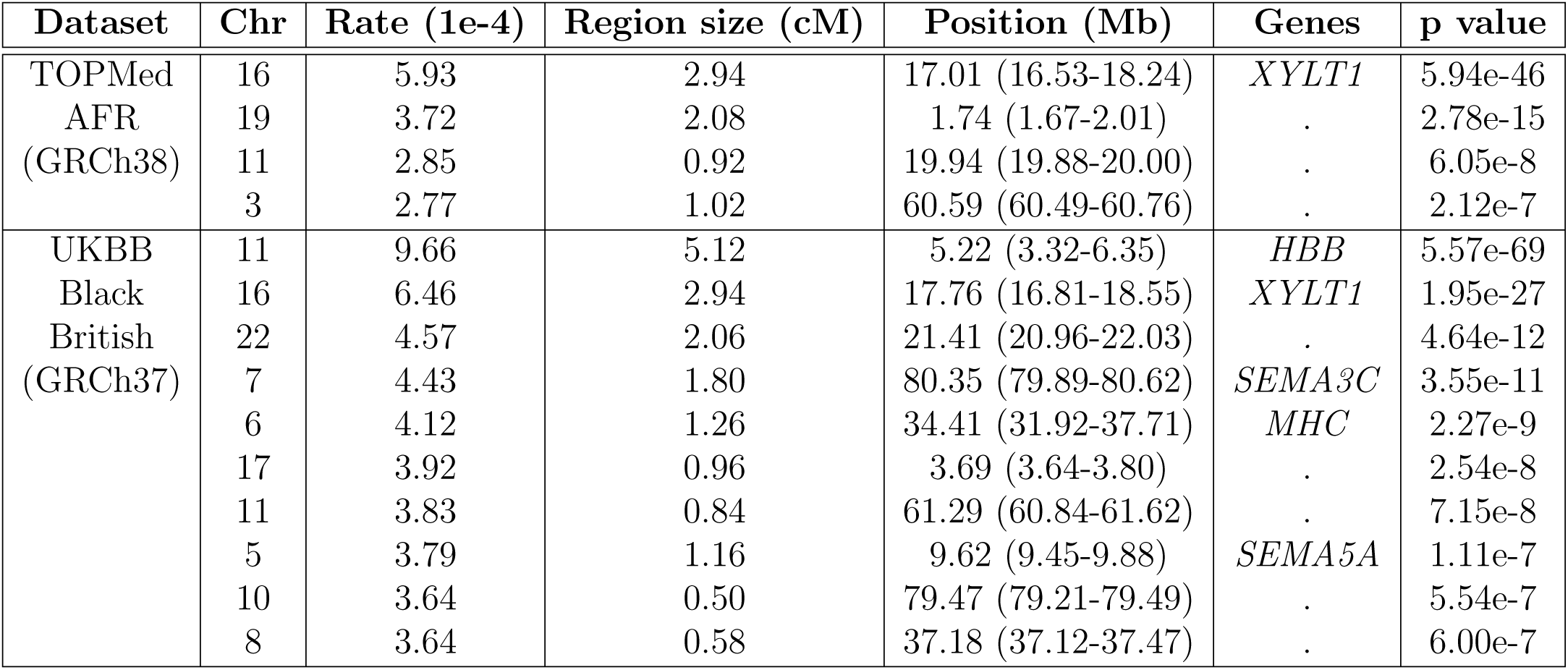
Loci detected in African ancestry selection scans. We report loci where identity-by-descent (IBD) rates exceed the multiple-testing analytical thresholds of 2.63e-4 and 3.55e-4 for the TOPMed AFR ancestry and UKBB Black British sample sets, respectively. The maximum IBD rate is given for each locus. Physical positions for the location of the maximum IBD rate and the span of excess IBD rates are shown in megabases (Mb). We report the size in centiMorgan (cM) of each region, which is defined to be a contiguous stretch of IBD rates exceeding the genome-wide significance threshold. Pedigree-based recombination maps from Halldorsson et al. [80] and Bhérer et al. [82] aligned to the GRCh38 and GRCh37 reference genomes are used for inferring IBD segments in the TOPMed and UKBB sample sets, respectively. p values are calculated assuming the null model that IBD rates are normally distributed. Annotated genes or gene complexes are discussed in the main text. The IBD segment detection threshold is 2.0 cM.

Using the original Temple et al. [14] selection scan workflow, twenty-four loci exceed the heuristic threshold of four standard deviations above the autosome-wide median in the ≥ 2.0 cM scan for the TOPMed AFR ancestry data [60]. Using our modified workflow with the multiple-testing corrections, only four of these twenty-four loci are genome-wide significant. Similarly, nineteen loci exceed our heuristic threshold of four standard deviations above the autosome-wide median in the ≥ 2.0 cM scan for the UKBB Black British data [60], only ten of which exceed our multiple-testing adjusted threshold.

Except for a 0.02 cM stretch of excess IBD rates in the *MHC* region, no loci are genome-wide significant in the TOPMed EUR2 ancestry data. The mean IBD rate is an order of magnitude larger for this group than for any other group. Recall that this European ancestry sample set is likely descendants from a small founder population. In such a demographic scenario, *de novo* sweeping alleles are more likely to be lost than in large populations.

Figure 3 shows the ≥ 3.0 cM IBD rates along the autosomes, the autosome-wide median, the heuristic four standard deviations above the median threshold, and the multiple-testing adjusted thresholds for the TOPMed EUR1, UKBB white British, UKBB Indian British, TOPMed AFR ancestry, and UKBB Black British groups. We report the statistically significant results of the ≥ 3.0 cM scan in Table S2.

### 3.5. Replicating selection signals in European ancestry groups

We previously reported eight of the eleven statistically significant loci in the TOPMed EUR1 scan selection scan [14]. The p value for the *LCT* gene (MIM: 603576) is so small that is cannot be represented in the 64-bit floating point system. The three loci not reported in our prior analysis of the TOPMed EUR1 ancestry data have been reported in other studies to be under selection. The genes *TLR1*, *TLR6*, and *TLR10* (MIM: 601194, 605403, and 606270) encode toll-like receptors that help initiate an immune response and may have been under selection in ancient Eurasians [38]. Gittelman et al. [83] have also suggested that an introgressed Neanderthal haplotype covering the *TLR1-6-10* genes may have been under selection. Multiple late cornified envelope (*LCE*) genes in the human epidermal complex are a few tens of kb from the significant locus on chromosome 1 and highly expressed in skin. The *HNF1B* gene (MIM: 189907) on chromosome band 17q12 is associated with diabetes and prostate cancer [84, 85].

Based on our simulation study of statistical power, we expect that hard sweeps from a single beneficial allele that are detected in the ≥ 3.0 cM scan will also be detected in the ≥ 2.0 cM scan. In the TOPMed EUR1 ancestry data, four significant loci in the ≥ 3.0 cM scan are also significant loci in the ≥ 2.0 cM scan. The signal near the *HNF1B* gene is barely genome-wide significant in ≥ 2.0 cM scan but is the third most significant in the ≥ 3.0 cM scan. The three loci significant in the ≥ 3.0 cM scan but not in the ≥ 2.0 cM scan are containing a family of keratin genes on chromosome 12 (*KRT*), a few hundred kb upstream of the immunoglobulin lambda genes (*IGL*), and in a gene-sparse region on chromosome band 16q12.3.

In the UKBB white British data, we observe ≥ 2.0 cM and ≥ 3.0 cM IBD rates exceeding our genome-wide significance threshold at many of the same loci significant in the TOPMed EUR1 ancestry analysis (Tables 2 and S2). Five of the twelve primary selection signals and none of the secondary selection signals in the Browning and Browning [24] analysis of the UKBB white British data are genome-wide significant in our scan. Two loci are genome-wide significant in the UKBB white British scan but not in the TOPMed EUR1 ancestry scan. The *CCR9* gene (MIM: 604738) encodes a chemokine receptor that plays an essential role in the mucosal immune system [86] and has been associated with increased COVID-19 outcome severity, especially in Europeans [87]. At this locus, Browning et al. [88] and Ding et al. [89] suggest that introgressed Neanderthal haplotypes may be selected for in South and East Asians, respectively. The *MAPT* gene (MIM: 157140) on chromosome band 17q21.31 is contained within a 900 kb polymorphic inversion that may have been subject to recent selection in European ancestry populations [90].

### 3.6. Shared selection signals across ancestry groups

In the UKBB Indian British data, we also observe excess ≥ 2.0 cM and ≥ 3.0 cM IBD rates at the *LCT*, *MHC*, and *TRPM1* regions (Tables 2 and S2). Romero et al. [91] suggest that northern European haplotypes carrying a putatively selected allele at *LCT* may be identical by descent to haplotypes in Indian pastoralists. Using the methods in Temple et al. [14], we infer an excess IBD outgroup comprising seventeen percent of the samples, which would be in the range of the selected allele frequency in Indian pastoralists in Romero et al. [91]. The rates of IBD alleles near the human leukocyte antigen (*HLA*) genes are known to be high in all HapMap populations [23], which is consistent with our selection scan results near the *HLA* genes. Excess IBD rates in the UKBB Indian British samples only overlap two of the three *HLA* regions reported to be under selection by Mathieson and Terhorst [37]. In contrast, excess IBD rates in the European ancestry samples overlap all three selected loci. The *TRPM1* gene (MIM: 603576) is a couple of Mb downstream of the *OCA2* gene (MIM: 203200), which has geographic patterns of population genetic variation indicative of strong selection [92]. Expression of *TRPM1* gene in melanocytes is positively correlated with melanin content and negatively correlated with melanoma [93]. Browning et al. [88] previously reported evidence of archaic selection around the *CCR9* gene in a South Asian ancestry group, but we do not observe a genome-wide significant signal of recent selection in our UKBB Indian British scan.

In the ≥ 2.0 cM scan for the UKBB Black British group and in the ≥ 3.0 cM scan for the TOPMed EUR1, UKBB white British, TOPMed AFR ancestry, and UKBB Black British groups, we observe a genome-wide significant locus on chromosome band 22q11.21. Contiguous stretches of excess IBD rates span between 2.06 to 5.56 cM in the different analyses, which is larger than many of the other genome-wide significant regions (Table 2, 3, and S2). The locations of maximum IBD rates are at roughly 21.50 Mb and 20.25 Mb between analyses using GRCh37 versus GRCh38 reference builds, which do not map to the same sets of genes. In their analysis of the UKBB white British data, Browning and Browning [24] reported that the selection signal is close to the *UBE2L3* gene (MIM: 603721), which is associated with multiple autoimmune diseases [94]. The *IGL* genes involved in the adaptive immune system are also a few hundred kb downstream of this region. Overall, there is no clear indication across analyses of which genes within this gene-dense region could explain this signal.

IBD rates spanning a couple of Mb on chromosome band 16p12.3 are genome-wide significant in the ≥ 2.0 cM scans for UKBB Indian British, TOPMed AFR ancestry, and UKBB Black British groups and in the ≥ 3.0 cM scans for TOPMed EUR1 ancestry and UKBB white British groups (Tables 2, 3, and S2). This region’s most extreme ≥ 2.0 cM IBD rates are 14.17 and 10.78 standard deviations above the autosome-wide means in the TOPMed AFR ancestry and UKBB Black British groups. This region’s maximum ≥ 2.0 cM IBD rate is only 6.05 standard deviations above the autosome-wide mean in the UKBB Indian British data. (For reference, the IBD rate at the *TRPM1* gene is 11.71 standard deviations above the autosome-wide mean in the TOPMed EUR1 ancestry group.) Excess IBD rates span at least 2.5 cM of this region in all of these analyses. Applying the subgroup anomaly detection method in Temple et al. [14] to the TOPMed AFR ancestry data, we fail to detect a singular excess IBD sharing cluster at this locus, which would have been indicative of a hard selective sweep.

This genomic region contains few genes, with the excess IBD rates entirely spanning the *XYLT1* gene (MIM: 608124). The *XYLT1* gene encodes the xylosyltransferase 1 enzyme, which initiates a chain reaction in the early maturation of skeletal cells. Linkage analysis in a consanguineous Turkish family associated a recessive missense mutation with a short stature syndrome [95]. Mutagenesis screening of mice also demonstrated disproportionate dwarfism from a recessive missense mutation in *XYLT1* [96]. This finding could thus be an example of recent selection across multiple continental ancestry groups that is not targeting immunity nor pigmentation-related genes.

### 3.7. African ancestry-specific recent selection signals

Some genome-wide significant loci are only found in the African ancestry analyses. For example, excess IBD rates also cover most of the *SEMA5A* gene (MIM: 609297) on chromosome 5. This gene encodes a protein specifically expressed around retinal axons in the optic nerve and helps maintain the axons’ structural integrity [97]. A deletion in the *SEMA5A* gene has been associated with autism spectrum disorders [98].

Around the genome-wide significant signal on chromosome band 7q21.11 in the UKBB Black British selection scans, we observe a subset of single nucleotide polymorphisms (GRCh37, chr7: 8,039,598; 80,624,286; 80,715,067) strongly differentiated between a group of excess IBD sharing and the rest of the sample [14]. These SNPs have frequencies between 72-79%, 15-20%, and 20-25% in the excess IBD sharing group, the rest of the sample, and the entire sample, respectively. The SNPs lie in the *SEMA3C* gene (OMIM: 602645). This gene encodes a protein involved in neuronal guidance. Expression of this gene is positively correlated with Wnt pathway activation, which is often dysregulated in brain tumor cancers [99]. We observe a genome-wide significant locus on chromosome band 11p15.4 in the ≥ 3.0 cM scan for TOPMed AFR ancestry samples and in the ≥ 2.0 and ≥ 3.0 cM scans for the UKBB Black British samples. This locus has more extreme IBD rates than the *XYLT1* gene in the UKBB Black British data. At this locus, we apply the Temple et al. [14] methods to the UKBB Black British data to detect a subset of single nucleotide polymorphisms (SNPs) strongly differentiated between a group of excess IBD sharing and the rest of the sample. We observe various well-differentiated SNPs (GRCh37, chr11: 5,221,233; 5,223,750; 5,214,301) within tens of kb of the hemoglobin beta gene (*HBB*, OMIM: 141900). These SNPs have frequencies between 81-85%, 14-19%, and 22-27% in the excess IBD sharing group, the rest of the sample, and the entire sample, respectively. Hemoglobins are proteins in red blood cells that transport oxygen to cells and tissues [100]. Mutations in the cluster of genes encoding the hemoglobin beta subunits are suspected to be targets of selection to reduce susceptibility to infections and malaria but also causes of sickle cell anemia and beta thalassemia disorders [101].

## 4. Discussion

In this paper, we model the correlation of detectable identity-by-descent segments along chromosomes to determine approximate genome-wide significance levels for an IBD rate-based selection scan. One of our approaches calculates the genome-wide significance level analytically, compared to permutation- and simulation-based approaches that are common in genetic studies but can be computationally intensive or intractable. Developing valid multiple-testing approaches is important for complex haplotype-based analyses instead of using the GWAS significance level of 5e-8, lest we inflate Type 1 errors or decrease the power to reject false null models. By properly accounting for correlations between test statistics, we can perform hypothesis tests finely spaced along the autosomes, thereby increasing statistical power.

Due to the speed of the msprime and tskibd methods for simulating IBD segments along entire chromosomes, we can measure the FWER in different demographic scenarios and under various experimental conditions. Many methods to detect recent selection have not measured the FWER in simulation studies, in large part because of the immense computation that would be involved, nor have they proposed multiple-testing corrections [20, 26, 32, 30, 17, 33, 34, 35, 31]. We find that our ≥ 2.0 and ≥ 3.0 cM scans have slightly anti-conservative and conservative control of the FWER, respectively. The asymptotic conditions of Temple and Thompson [59] are less valid in the ≥ 2.0 cM scan, which may explain its anti-conservative behavior. The asymptotic conditions of Temple and Thompson [59] are more reasonable in the ≥ 3.0 cM scan, but the Siegmund and Yakir [54] analytical approximation is conservative for true OU processes.

Unless the genetic data has low coverage or poor genotyping quality such that detecting IBD segments less than 3.0 cM is inaccurate [102, 103], we recommend using the anticonservative ≥ 2.0 cM scan over the conservative ≥ 3.0 cM scan, which has limited power. The ≥ 3.0 cM scan has limited power to detect hard sweeps of *s <* 0.015, which Schrider and Kern [43] describe as strong selection. On the other hand, we find that the ≥ 2.0 cM scan has some power when *s* ≤ 0.010 and considerable power when *s >* 0.01. Indeed, the heuristic threshold of Temple et al. [14] corresponds to the expected IBD rate of an *s* = 0.017 sweep in the TOPMed EUR1 ancestry samples. Some methods claim to have the power to detect sweeps

where *s <* 0.010 [39, 40, 34, 35]. However, these methods do not address multiple testing. We suggest that selection coefficients *s <* 0.01 and *s* ≥ 0.010 may describe undetectable and detectable recent sweeps once multiple testing is accounted for.

We consider the hard sweep model in our power simulations, which is one of many alternative models that could explain excess IBD rates. The pairwise IBD rate test does not resolve the classification of hard and soft sweeps versus recurrent sweeps versus balancing selection versus other mechanisms, which is a topic of growing interest in the field [104, 40, 43, 105]. We observe that hard sweeps detected in the ≥ 3.0 cM scan are almost always detected in the ≥ 2.0 cM scan, in which case loci significant in the ≥ 3.0 cM scan but not in the ≥ 2.0 cM scan may not be the result of a hard sweep. In practice, we should account for the fact that conducting scans with multiple different segment length thresholds is another form of multiple testing (Appendix A.2). Temple et al. [14] also propose various diagnostics as characteristic of a hard sweep, particularly that of a single majority haplotype cluster with excess IBD rates and a reduction in the diversity of common variants.

Failing to adjust for multiple testing properly can be cause for concern in discovery studies. In our study, we investigate signals of natural selection in human populations, in which significant findings could be misinterpreted or misappropriated [106]. After adjusting for multiple testing, we identify eleven or fewer statistically significant results in any given ancestry or ethnicity cohort. In contrast, Akbari et al. [107] report more than three hundred independent significant results of recent selection, using a novel rescaling to address genomic inflation. We have validated control of the FWER in simulation studies, whereas Akbari et al. [107] have not.

We find that the four standard deviations above the autosome-wide median threshold used in our previous work [14] is nearly identical to our new multipletesting corrections for the TOPMed EUR1 ancestry samples but that the heuristic IBD rate threshold is not large enough for studies on other ancestry groups. In African ancestry samples, we suggest that the many loci with IBD rates exceeding the heuristic threshold may be false positives. Nevertheless, in these African ancestry samples, we observe excess IBD rates around the *XYLT1* gene on the same relative magnitude as those around pigmentation genes believed to be under selection in European ancestry samples. This result indicates possible selection on skeletal cell development, whereas genes implicated in many prior selection studies are involved in immunity and pigmentation.

Replicating genome-wide significant results in different datasets and using different parameter configurations helps validate scientific results. Around many significant loci we use an automated workflow to show excess IBD rates in datasets of similar ancestry compositions but with different reference genomes and sequencing technologies. We find that IBD rates around the *XYLT1* gene are genome-wide significant in European ancestry, African ancestry, and the UKBB Indian British groups. This pattern of putatively recent selection in multiple ancestry groups is also present at *MHC*, an immunity gene complex broadly believed to be under some form of balancing selection [23]. Running our automated scan for recent selection in other European and African ancestry datasets or in other ancestry groups could corroborate our results and/or existing selection studies, for instance, selection at the *FADS* genes (MIM: 606148, 606149) [108, 38], and the *EDAR* gene (MIM: 604095) [109, 34].

Two limitations of our selection scan are genome size and sample size. To reliably estimate autosome-wide mean and standard deviations and the exponential decay parameter, we require more than 400 cM of genetic data. Additionally, the IBD rates along the chromosomes should not be zero, which happens when the sample size is too small to observe IBD segments ≥ 2.0 cM. For human genetics studies, one thousand samples is likely sufficient to apply our methodology [14, 60], albeit we recommend the analysis of at least a few thousand samples when such is available. In the case of small samples, one can review the scan plots output from the automated workflow to assess if the IBD rates are truncated to zero.

Analyzing chromosome 2 for the 1737 whole genome sequences in the TOPMed African ancestry data takes less than half a day with 8 CPUs, and analyzing chromosome 2 for 2500 Indian British samples in the UKBB SNP array data takes less than thirty minutes with 8 CPUs. Temple [60] shows that the ≥ 2.0 cM selection scan for two thousand randomly selected samples from the UKBB white British 410k data provides similar results to our analysis of the entire dataset. Compared to GWAS, where more samples leads to a smaller standard error and thereby more power to detect a nonzero regression effect, our selection scan is a test of neutrality for a stochastic process. It only requires enough samples such that the IBD rates along the chromosomes are not zero. Using more samples than necessary can lead to substantial runtime, random access memory (RAM), and disk memory costs: analyzing chromosome 2 for all UKBB white British samples takes nearly a week with 16 CPUs and 256 GB RAM, and the analysis of all autosomes leaves a memory footprint of 2.8 TB.

The hypothesis test and our multiple-testing corrections are so far limited to analyzing the autosomes of samples from large populations with *panmixia*. Skov et al. [110] report fourteen regions with extended common haplotypes as possible examples of strong archaic selection on the human X chromosome. Our multiple-testing corrections still apply to the X chromosome but would require separate estimation of the baseline IBD rate and the correlation parameter, which could be noisy when using data from only one chromosome (Figure S1). We restrict our analyses of admixed samples to those subsets with a large majority of one ancestry class. Selection studies in Native American populations have consisted of small sample sizes [108] relative to our study. Many admixed samples from TOPMed and other data have considerable but still minority compositions of Native American ancestry [111, 112, 57]. Future work in admixed samples could consider summary statistics and correlations of haplotype segments that are both detectably IBD and from the same ancestry group. Finally, our modeling assumptions are unreasonable in samples from a small population where the upper tail probabilities of high IBD rates can be greater than those of normal distributions. Modeling higher variance processes, like a Ĺevy-driven OU process [113], may be necessary to control the FWER of our selection scan when studying samples from founder or domesticated populations.

## Data and code availability

The methodology is implemented in the https://github.com/sdtemple/isweep Python package as a module, which is available under the CC0 1.0 Universal License. Scripts to conduct the simulation studies are available under the v1.0 tag at https://github.com/sdtemple/isweep/papers/mult-test-paper/.

## Acknowledgments

This research has received funding from the US National Human Genome Research Institute of the National Institutes of Health under award number HG005701. S.D.T. also acknowledges funding support from the US Department of Defense National Defense Science and Engineering Graduate Fellowship, the US National Institutes of Health T32 GM081062 Predoctoral Training Grant in Statistical Genetics, and Schmidt Sciences, LLC. This research has used the UK Biobank Resource under Application Number 19934. Molecular data for the Trans-Omics in Precision Medicine (TOPMed) program was supported by the National Heart, Lung, and Blood Institute (NHLBI). The content of this article is solely the responsibility of the authors and does not necessarily represent the official views of the National Institutes of Health. Core support, including centralized genomic-read mapping and genotype calling along with variant quality metrics and filtering, was provided by the TOPMed Informatics Research Center (3R01HL-117626-02S1; contract HHSN268201800002I). Core support, including phenotype harmonization, data management, sample-identity QC, and general program coordination, was provided by the TOPMed Data Coordinating Center (R01HL-120393; U01HL-120393; contract HHSN268201800001I). See supplemental information for acknowledgments of individual studies in the TOPMed data. We thank Ruoyi Cai for helpful discussions about UKBB, Kelsey Grinde for helpful discussions about the Ornstein-Uhlenbeck process, and Elizabeth Thompson, Kelley Harris, and Ryan Waples for feedback on early drafts of this manuscript.

## Author contributions

S.D.T. planned the study, wrote the software, conducted the analysis, and wrote the manuscript. S.D.T and S.R.B. developed the method. S.R.B. proposed the study and contributed to editing the manuscript.

## Web resources

- https://github.com/browning-lab/hap-ibd: detecting identity-by-descent segments
- https://github.com/browning-lab/ibd-ends: detecting identity-by-descent segments
- https://tskit.dev/: utilities for tree sequences, including msprime
- https://github.com/bguo068/tskibd: deriving identity-by-descent segments from tree sequences
- deCODE [80] genetic map: https://www.science.org/doi/suppl/10.1126/science.aau1043/suppl file/aau1043 datas3.gz
- Bhérer et al. [82] genetic map: https://github.com/cbherer/Bherer etal SexualDimorphismRecombination
- UCSC Genome Browser https://genome.ucsc.edu

## Appendix

### A.1. Accuracy of identity-by-descent segment detection

In a pilot study of ten population bottleneck simulations, we place mutations on the msprime tree sequence at a genome-wide rate of 1e-8. Then, we infer IBD segments with the hap-ibd and ibd-ends analysis workflow in Temple et al. [14]. Figures S10A-B illustrate one simulation of the true tskibd and inferred IBD rate processes across the genome. We observe similar genome-wide median IBD rates and significance thresholds between the true and inferred IBD rate processes. The inferred IBD rates are within 95 to 105% of the corresponding true IBD rates (Figure S10C). Across the ten simulations, the average estimates of *θ*^^^ are 69 and 75, and the average standard deviations *σ*^_1:_*_M_* are 19 and 20 for the true and inferred IBD rate processes, respectively.

We also conduct the same pilot study for five simulations of the constant size population of fifty thousand diploid individuals. Figure S11A illustrates one simulation of the inferred IBD rates divided by the true IBD rates across the genome. The inferred IBD rates are within 90 to 95% of the corresponding true IBD rates. We also observe a pattern of higher inferred IBD rates near the chromosome ends than the genome-wide median IBD rate. We run ibd-ends again with the hidden parameter ne=50000, observing only marginal differences compared to the software default setting (Figures S11B-C). For this demographic scenario, the differential detection accuracy of ibd-ends near chromosome ends could affect the control of the FWER.

### A.2. Multiple testing by using different segment length thresholds

A multiple-testing adjustment in a joint ≥ 2.0 and ≥ 3.0 cM scan should not be drastically different from the multiple-testing adjustment in the ≥ 2.0 cM scan because the individual ≥ 2.0 and ≥ 3.0 cM scans are highly correlated. In the population bottleneck simulations, we calculate that the medians of estimates *θ*^^^ for the ≥ 2.0 and ≥ 3.0 cM IBD rates are roughly 63 and 40. We also calculate that the median of crosscorrelations between the ≥ 2.0 and ≥ 3.0 cM (standardized) IBD rates is roughly 0.68. Next, we simulate a 2-dimensional standardized OU process two thousand times with the crosscorrelation parameter *ρ* = 0.68 and autocorrelation parameters *θ*_1_ = 63 and *θ*_2_ = 40. The data for each simulation is equivalent to 10 chromosomes of length 100 cM and testing every 0.02 cM. From these simulations, we calculate that the 95th percentiles of the maxima of marginal OU processes with *θ*_1_ = 63 and *θ*_2_ = 40 are 4.36 and 4.24, which correspond to genome-wide significance levels of 6.50e-6 and 1.12e-5. We also calculate that the 95th percentile of the maximum of the 2-dimensional OU process is roughly 4.47, corresponding to a significance level of 3.91e-6.

## Supplementary figures

**Figure S1:**
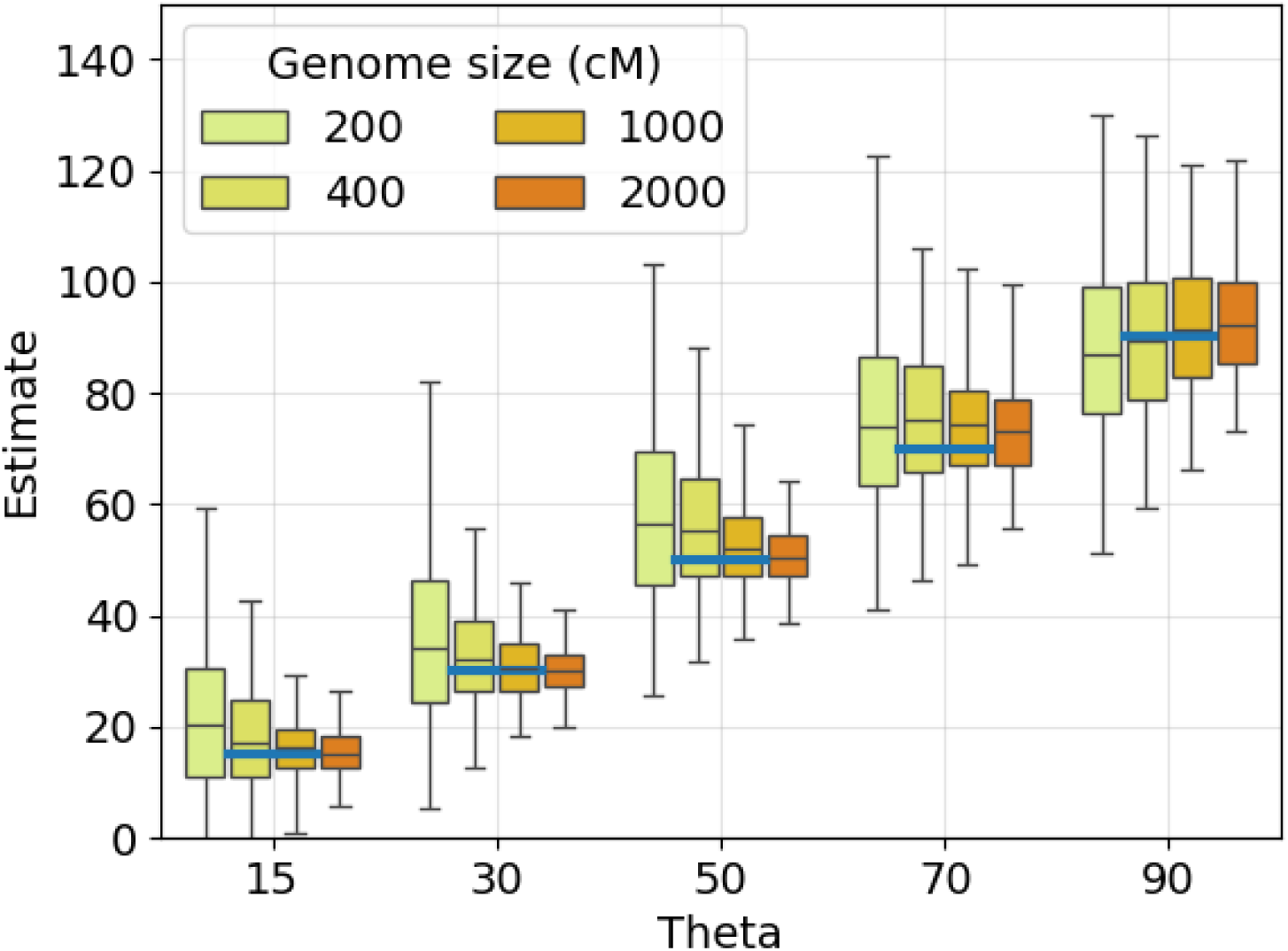
Estimating the exponential decay parameter *θ* from simulated Ornstein-Uhlenbeck processes. The 1st, 25th, 50th, 75th, and 99th percentiles of estimates *θ*^^^ (y-axis) are shown for true *θ* (x-axis and horizontal blue lines). We estimate *θ* with different genome lengths (colors in legend) and step size 0.02 cM. Percentiles are taken over five hundred simulations for each *θ*.

**Figure S2:**
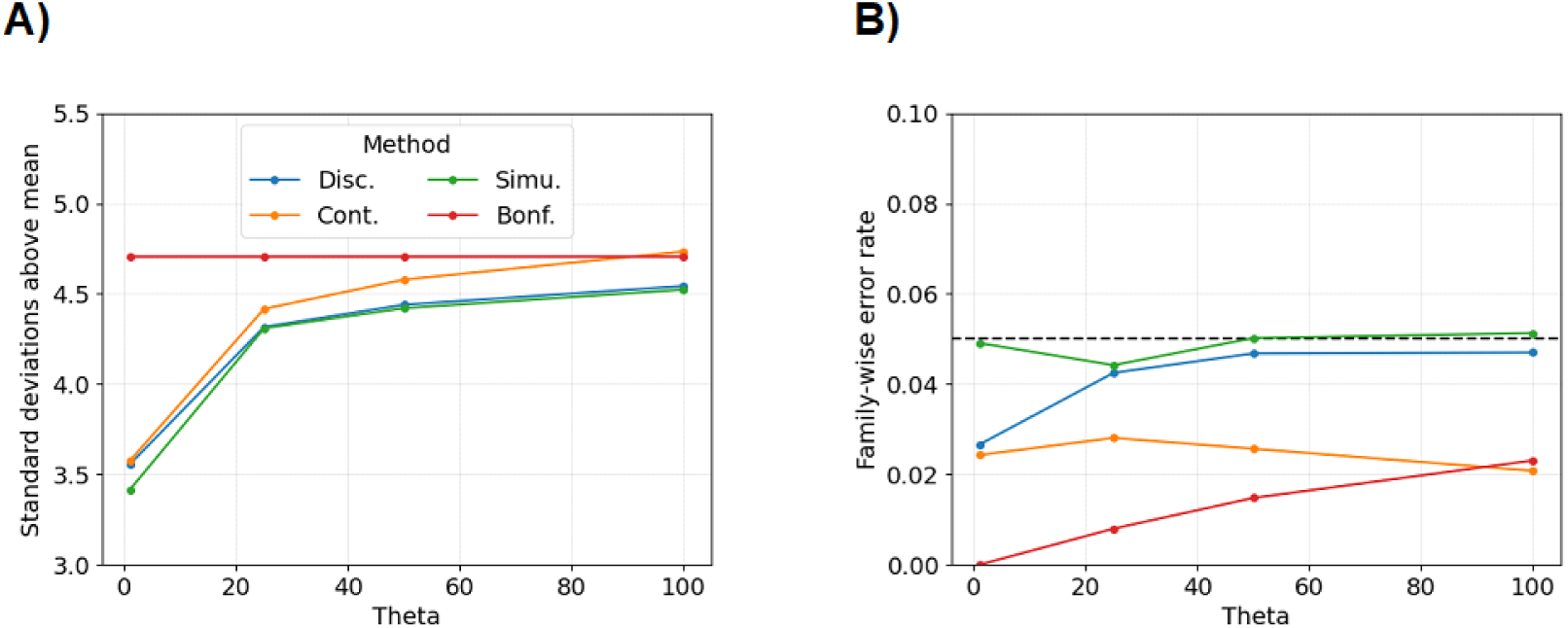
Multiple-testing approaches in simulations of Ornstein-Uhlenbeck processes. Line plots show A) standard deviations above the mean thresholds or B) family-wise error rates (y-axis) with different *θ* (x-axis). The multiple-testing approaches are (blue) the discrete-spacing analytical approximation, (orange) the continuous-spacing analytical approximation, (green) the simulation-based approach, and (red) the Bonferroni correction. The simulation-based approach is based on ten thousand simulations. The step size is hypothesis testing every 0.05 cM (50 kb). The data for each simulation is equivalent to twenty chromosomes, each of length 100 cM.

**Figure S3:**
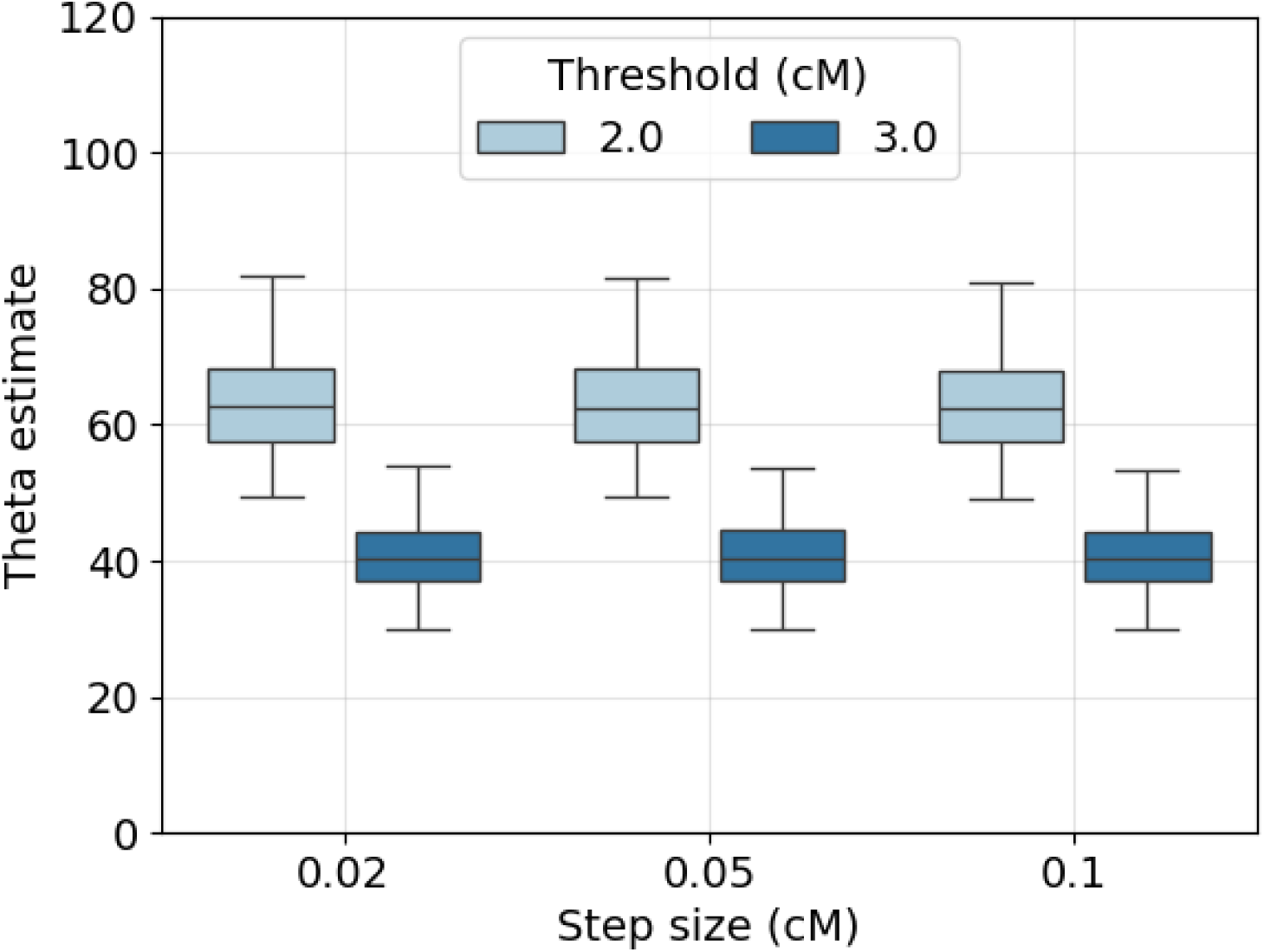
Estimating the exponential decay parameter *θ* from simulated IBD rate processes with different cM length thresholds. Box plots show the 1st, 25th, 50th, 75th, and 99th percentiles of estimates *θ*^^^ using the IBD rate processes with simulated true IBD segments (dark blue) ≥ 2.0 cM and (light blue) ≥ 3.0 cM from tskibd. Estimates *θ*^^^ are based on autocovariances calculated at different step sizes (x-axis). There are fifteen hundred simulations for each step size. The demographic model is the population bottleneck. The data for each simulation is equivalent to ten chromosomes of uniform length 100 cM.

**Figure S4:**
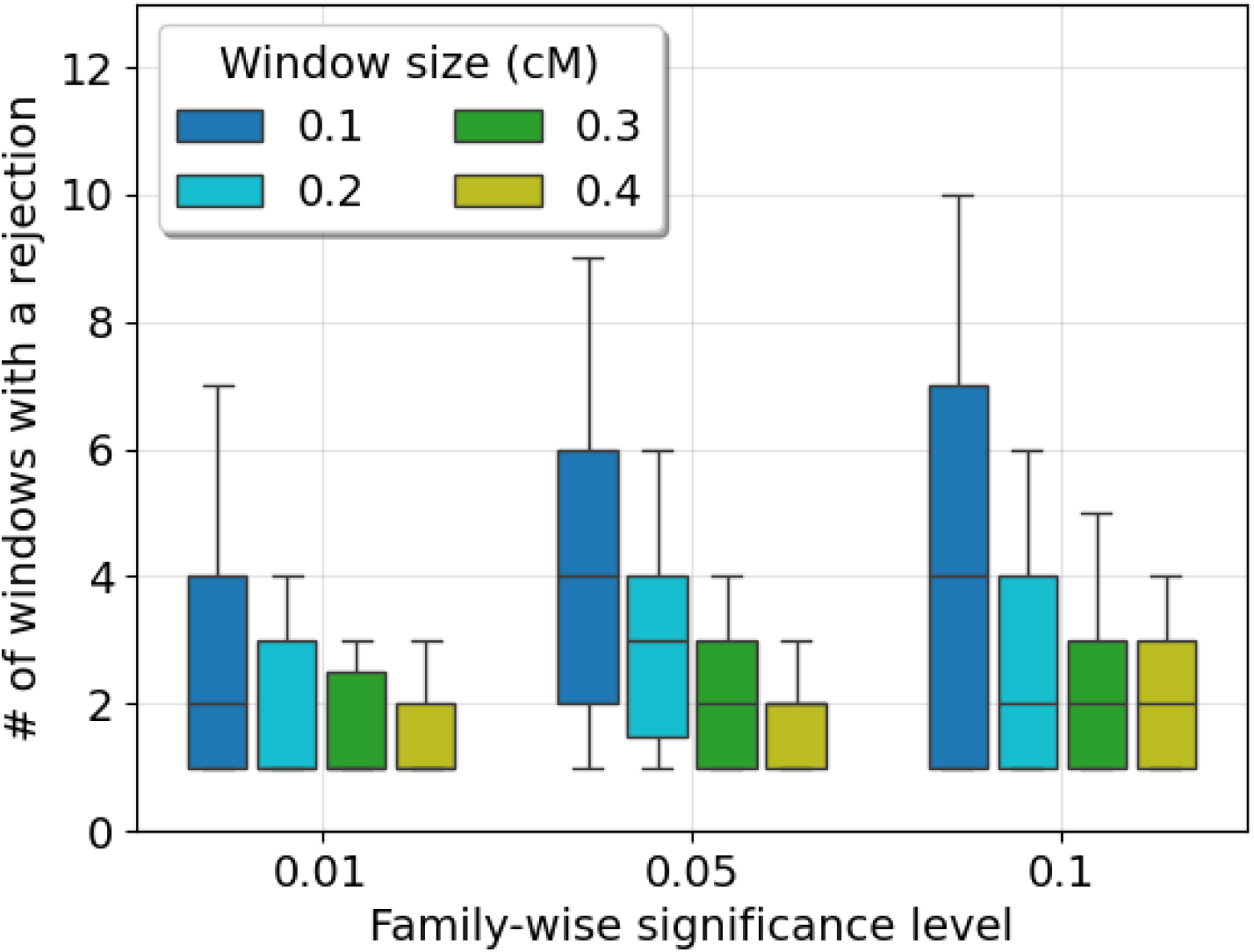
The number of windows with a rejected hypothesis test. Box plots show the 10th, 25th, 50th, 75th, and 90th percentiles of the number of non-overlapping windows with at least one rejection of the null hypothesis (y-axis). Windows sizes are 0.1, 0.2, 0.3, and 0.4 cM (colors in legend) with IBD rates calculated every 0.02 cM. Simulations in which there are no genome-wide significant tests are not included in the box plots. The multiple-testing method is the discrete-spacing analytical approximation using true IBD segments ≥ 2.0 cM. There are five hundred simulations for each family-wise significance level (x-axis). The demographic model is the population bottleneck. The data for each simulation is equivalent to ten chromosomes of uniform length 100 cM.

**Figure S5:**
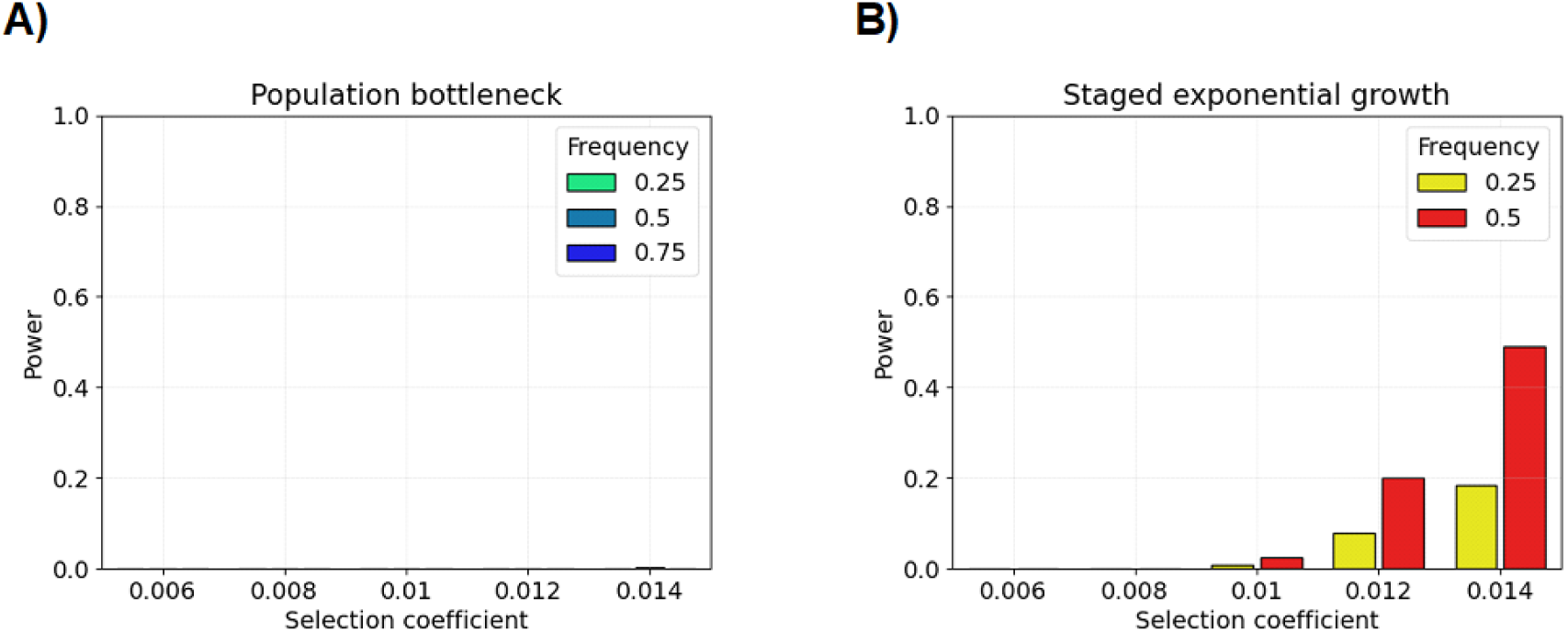
Power simulations for the ≥ 3.0 cM scan in different demographic models. Bar plots show the statistical power (y-axis) in the A) population bottleneck and B) staged exponential growth models using true IBD segments ≥ 2.0 cM overlapping the selected allele. Power is the proportion of tests where the null model is rejected at the p value threshold corresponding to the 0.05 family-wise significance level. Hypothesis testing is based on the discrete-spacing analytical threshold using the step size 0.02 cM. There are two hundred simulations for each pair of selection coefficient (x-axis) and current-day allele frequency (colors in legend).

**Figure S6:**
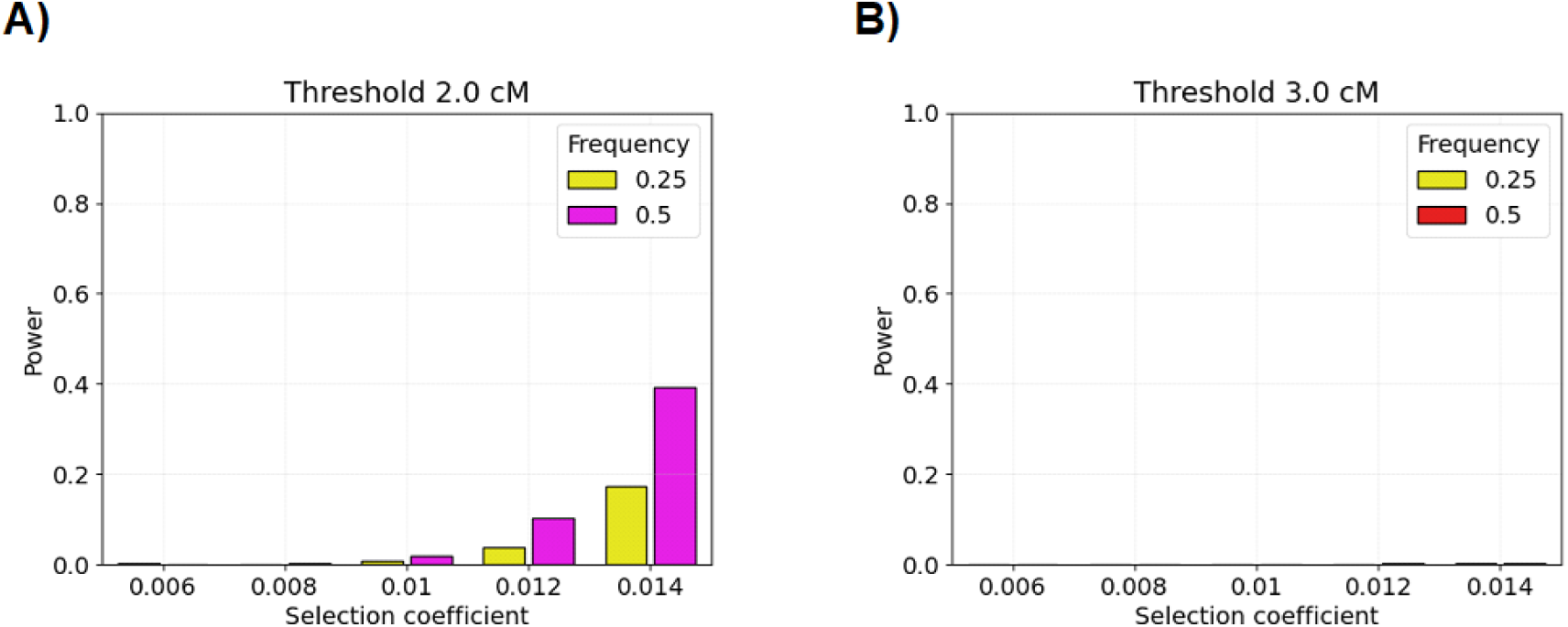
Power simulations in a constant size population. Bar plots show the statistical power (y-axis) using true IBD segments A) ≥ 2.0 cM or B) ≥ 3.0 cM overlapping the selected allele. Power is the proportion of tests where the null model is rejected at the p value threshold corresponding to the 0.05 family-wise significance level. Hypothesis testing is based on the discrete-spacing analytical threshold using the step size 0.02 cM. There are two hundred simulations for each pair of selection coefficient (x-axis) and current-day allele frequency (colors in legend).

**Figure S7:**
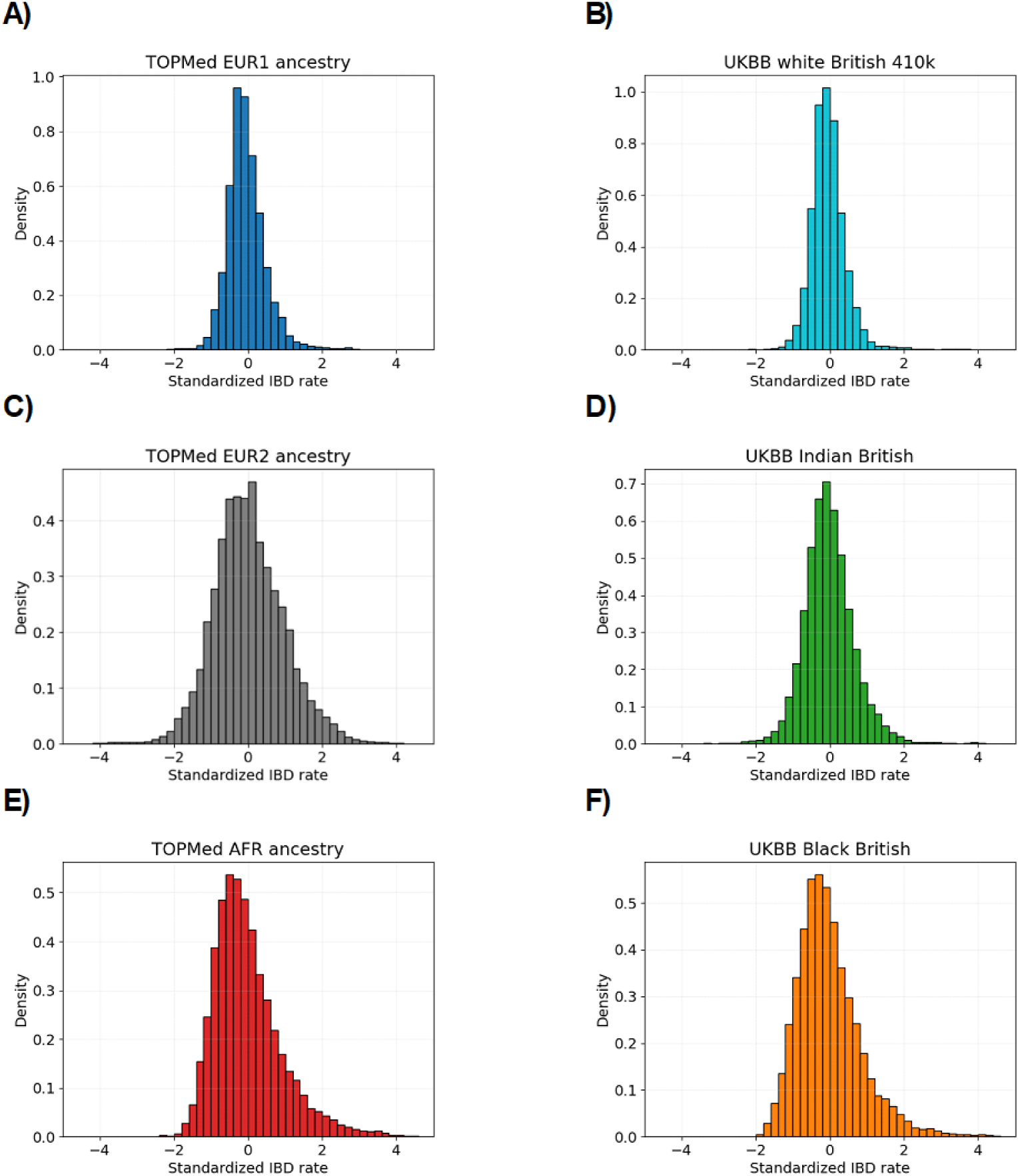
Histograms of IBD rates in human populations. The standardized IBD rates ≥ 2.0 cM (*x*-axis) are shown for A) TOPMed EUR1, B) UKBB white British, C) TOPMed EUR2, D) UKBB Indian British, E) TOPMed AFR ancestry, and F) UKBB Black British sample sets. Each histogram has fifty bins, and the x-axes range from −5 to 5.

**Figure S8:**
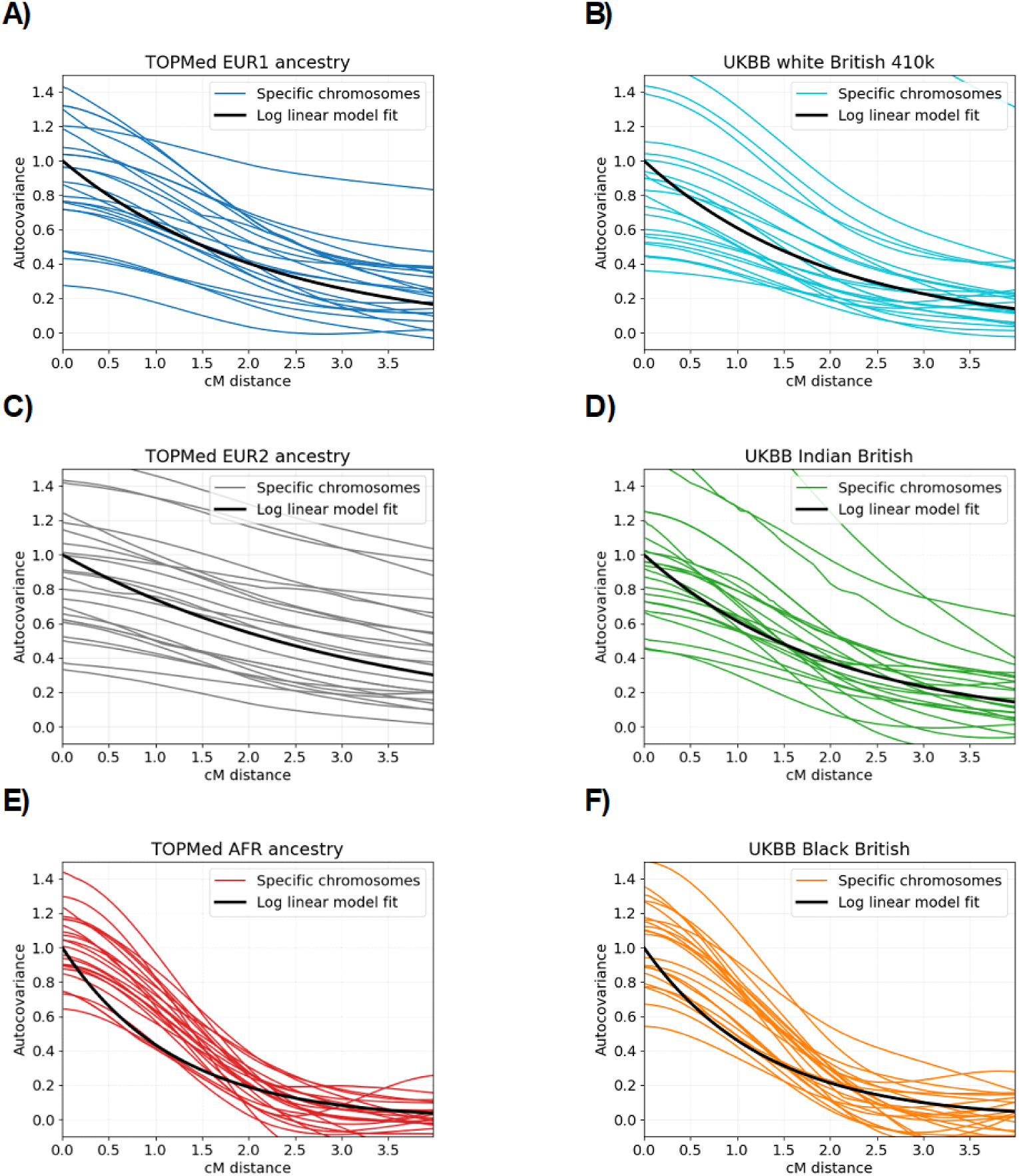
Estimating exponential decay parameter *θ* in real data. Each faint colored line shows estimated autocovariances (y-axis) for different cM distances (x-axis) and a specific chromosome. The black lines show the predicted autocovariances from the fitted Ornstein-Uhlenbeck processes using estimates *θ*^^^. The data for each subplot is based on A) TOPMed EUR1 ancestry, B) UKBB white British, C) TOPMed EUR2 ancestry, D) UKBB Indian British, E) TOPMed AFR ancestry, and F) UKBB Black British sample sets.

**Figure S9:**
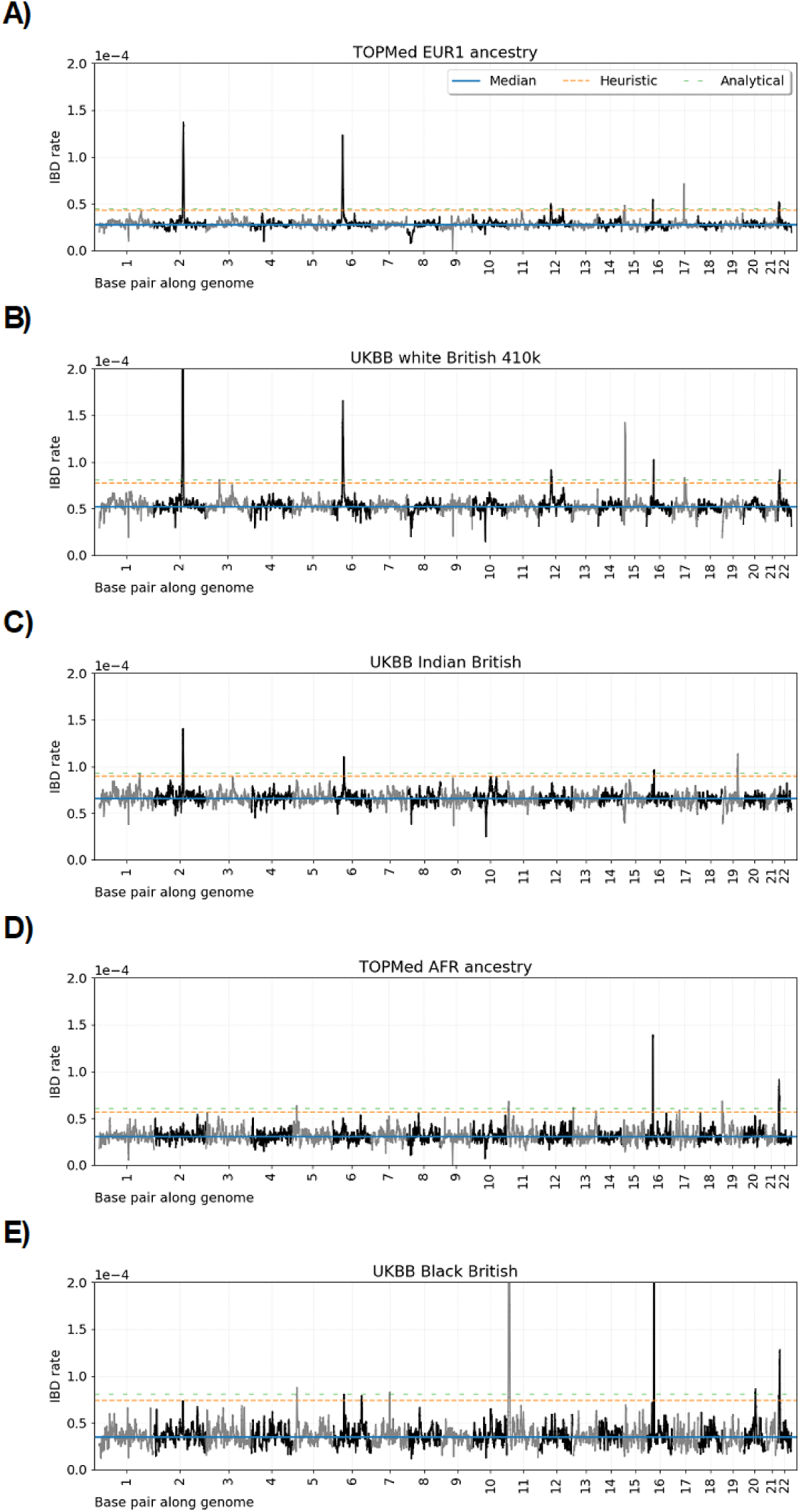
Genome-wide IBD rate scans using the ≥ 3.0 cM threshold. Line plots show IBD rates every 0.02 cM (y-axis) for base pair positions along twenty-two human autosomes. The data for each subplot is based on A) TOPMed EUR1 ancestry, B) UKBB white British, C) UKBB Indian British, D) TOPMed AFR ancestry, and E) UKBB Black British sample sets. Horizontal dashed lines show (blue) the genome-wide median IBD rate, (orange) the heuristic threshold of four standard deviations above the median, and (green) the analytical multiple-testing threshold.

**Figure S10:**
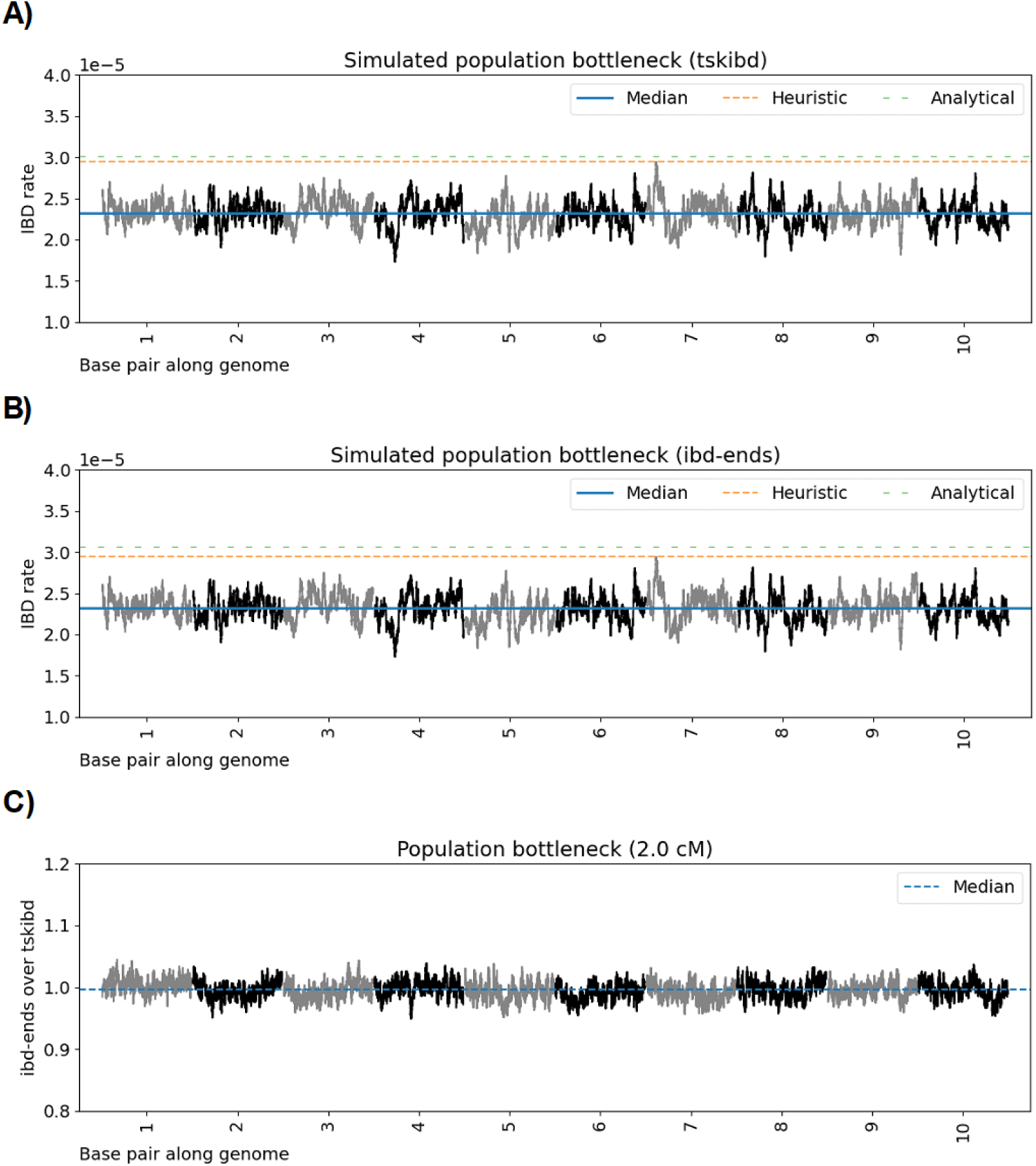
Genome-wide IBD rate scan in a simulated population bottleneck scenario. Line plots show ≥ 2.0 cM IBD rates (y-axis) for cM positions along ten simulated chromosomes. Scans are based on A) tskibd true IBD segments [6] or B) ibd-ends inferred IBD segments [24]. In C), we divide the IBD rates in B) from those in A). Each chromosome is 100 cM. The IBD rate is calculated every 0.02 cM. Data is based on twenty-five hundred diploid samples from the simulated population bottleneck demographic scenario. Horizontal dashed lines show (blue) the genome-wide median IBD rate, (orange) the heuristic threshold of four standard deviations above the median, and (green) the discrete-spacing analytical threshold). The family-wise significance level is 0.05.

**Figure S11:**
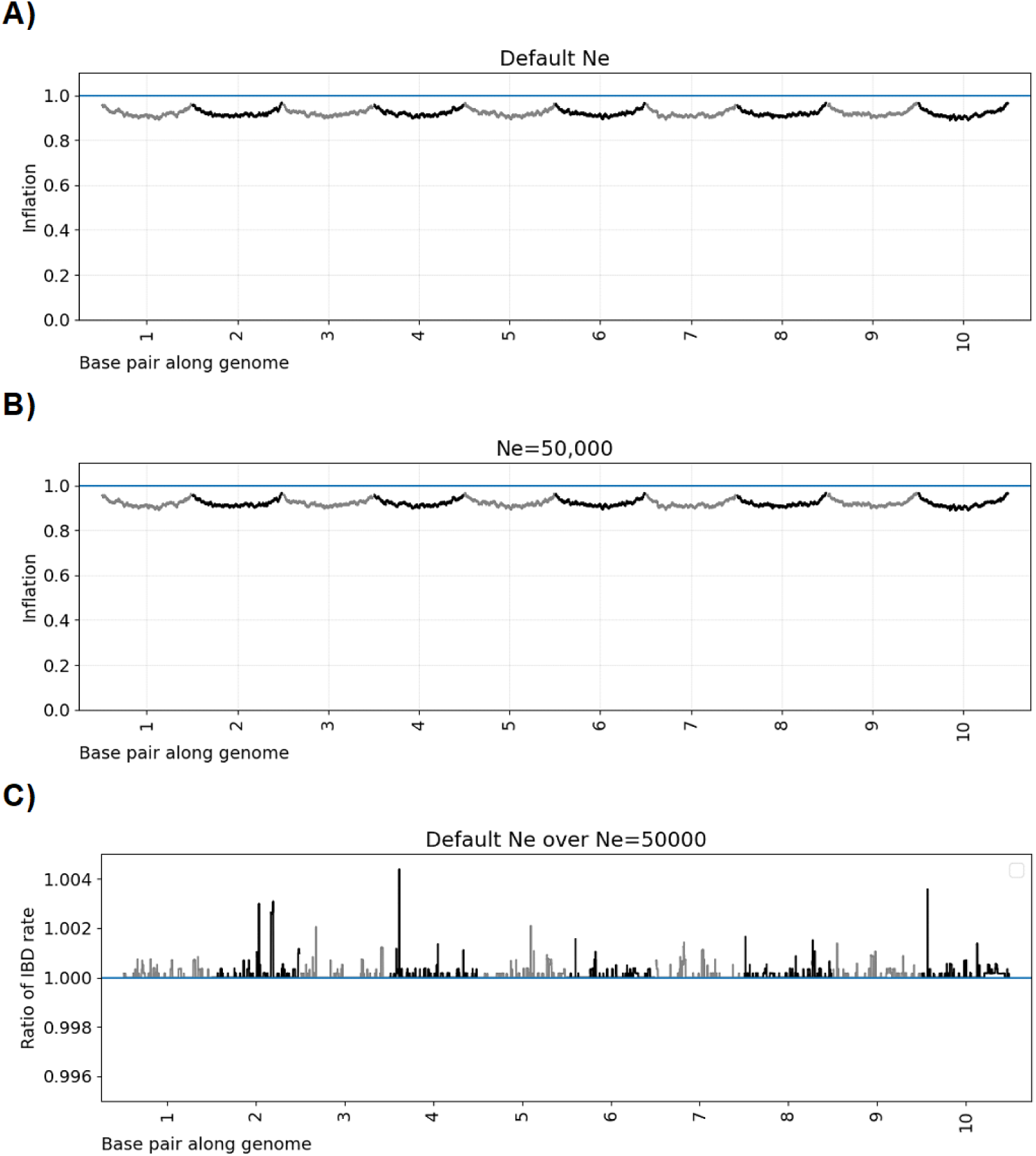
Genome-wide IBD rate scan in a simulated constant population size scenario. Line plots show inferred IBD rates over true IBD rates (y-axis) for cM positions along ten simulated chromosomes. Scans are based on using ibd-ends’s A) default ne setting versus B) ne=50000. In C), we divide the inferred IBD rates in A) and B). Each chromosome is 100 cM. The IBD rate is calculated every 0.02 cM. Data is based on twenty-five hundred diploid samples from the simulated scenario of a constant population of fifty thousand individuals. The segment detection threshold is ≥ 2.0 cM.

## Supplementary tables

**Table S1:** Significance levels and family-wise error rates after multiple-testing corrections with IBD segments ≥ 3.0 cM. Significance levels are adjusted for multiple testing based on scans over 10 chromosomes of size 100 cM and tests every 0.02 cM (50,000 total tests). The multiple-testing analytical and simulation-based thresholds are based on a fitted Ornstein-Uhlenbeck process. Each simulation has a different threshold as a result of estimating *θ*. Family-wise error rate (FWER) is the percentage of five hundred genome-wide scans with at least one statistically significant result. The demographic scenario is the population bottleneck.

| **Family-wise**<br>**level** | **Adjusted**<br>**Analytical** | **Simulation** | **Bonferroni** | **FWER**<br>**Analytical** | **Simulation** |
| --- | --- | --- | --- | --- | --- |
| 0.01 | 1.58e-6 | 2.01e-6 | 2.08e-7 | 0.008 | 0.012 |
| 0.05 | 9.23e-6 | 1.06e-5 | 1.04e-6 | 0.030 | 0.034 |
| 0.10 | 2.03e-5 | 2.29e-5 | 2.08e-6 | 0.066 | 0.078 |

**Table S2:** Loci detected in the ≥ 3.0 cM selection scans. We report loci where identity-by-descent (IBD) rates exceed the discrete-spacing analytical thresholds of 0.45e-4, 0.81e-4, 0.93e-4, 0.61e-4, and 0.81e-4 for the TOPMed EUR1 ancestry, UKBB white British, UKBB Indian British, TOPMed AFR ancestry, and UKBB Black British sample sets, respectively. The maximum IBD rate is given for each locus. Physical positions for the location of the maximum IBD rate and the span of excess IBD rates are shown in megabases (Mb). We report the size in centiMorgan (cM) of each region, which is defined to be a contiguous stretch of IBD rates exceeding the genome-wide significance threshold. Pedigree-based recombination maps from Halldorsson et al. [80] and Bhérer et al. [82] aligned to the GRCh38 and GRCh37 reference genomes are used for inferring IBD segments in the TOPMed and UKBB sample sets, respectively. p values are calculated assuming the null model that IBD rates are normally distributed. Annotated genes or gene complexes are discussed in the main text.

| Dataset | Chr | Rate (1e-4) | Region size (cM) | Position (Mb) | Genes | p value |
| --- | --- | --- | --- | --- | --- | --- |
| TOPMed<br>EUR1<br>(GRCh38) | 2 | 1.37 | 7.94 | 134.84 (132.29-140.09) | *LCT* | 2.59e-187 |
|  | 6 | 1.23 | 8.04 | 31.03 (23.91-36.38) | *MHC* | 2.14e-143 |
|  | 17 | 0.71 | 3.44 | 37.68 (36.33-38.44) | *HNF1B* | 2.59e-31 |
|  | 16 | 0.54 | 3.00 | 17.74 (16.73-18.72) | *XYLT1* | 8.50e-13 |
|  | 22 | 0.52 | 5.12 | 20.23 (19.15-21.10) | . | 6.37e-11 |
|  | 12 | 0.50 | 5.10 | 51.38 (48.80-53.20) | *KRT* | 1.74e-9 |
|  | 15 | 0.48 | 2.28 | 31.30 (30.46-32.15) | *TRPM1* | 4.19e-8 |
| UKBB<br>white<br>British<br>410k<br>(GRCh37) | 2 | 3.98 | 8.72 | 135.91 (132.86-141.34) | *LCT* | underflow |
|  | 6 | 1.65 | 7.92 | 30.80 (23.91-36.34) | *MHC* | 2.63e-74 |
|  | 15 | 1.42 | 3.50 | 30.94 (30.16-32.92) | *TRPM1* | 1.26e-47 |
|  | 16 | 1.02 | 3.28 | 18.25 (16.37-18.95) | *XYLT1* | 6.40e-16 |
|  | 22 | 0.91 | 3.46 | 21.53 (20.98-21.61) | . | 1.91e-10 |
|  | 12 | 0.91 | 5.12 | 51.78 (49.38-53.76) | *KRT* | 2.04e-10 |
|  | 17 | 0.83 | 1.26 | 36.18 (35.44-36.49) | *HNF1B* | 4.16e-7 |
| UKBB<br>Indian<br>British<br>(GRCh37) | 2 | 1.41 | 5.12 | 136.97 (134.36-139.51) | *LCT* | 1.69e-36 |
|  | 19 | 1.12 | 4.92 | 50.23 (48.47-50.74) | . | 5.25e-16 |
|  | 6 | 1.10 | 3.12 | 33.96 (33.02-36.34) | *MHC* | 4.70e-14 |
|  | 16 | 0.96 | 2.80 | 18.06 (16.83-18.28) | *XYLT1* | 1.65e-7 |
| TOPMed<br>AFR<br>(GRCh38) | 16 | 1.39 | 3.44 | 17.73 (16.45-19.09) | *XYLT1* | 1.31e-63 |
|  | 22 | 0.92 | 5.56 | 20.26 (18.95-21.10) | . | 3.30e-21 |
|  | 19 | 0.69 | 1.98 | 1.78 (1.72-2.10) | . | 3.56e-9 |
|  | 11 | 0.68 | 2.74 | 5.23 (3.83-5.75) | *HBB* | 4.83e-9 |
|  | 5 | 0.37 | 1.56 | 9.44 (9.20-9.72) | *SEMA5A* | 2.34e-7 |
| UKBB<br>Black<br>British<br>(GRCh37) | 11 | 3.79 | 7.78 | 4.72 (2.76-6.92) | *HBB* | 2.78e-271 |
|  | 16 | 2.52 | 4.14 | 17.40 (16.06-19.14) | *XYLT1* | 4.12e-109 |
|  | 22 | 1.28 | 4.74 | 21.54 (19.64-22.33) | . | 1.68e-21 |
|  | 5 | 0.88 | 1.64 | 9.62 (9.34-9.90) | *SEMA5A* | 5.77e-8 |
|  | 20 | 0.86 | 1.54 | 40.93 (39.47-40.99) | . | 1.30e-7 |
|  | 7 | 0.83 | 0.50 | 80.35 (80.08-80.40) | *SEMA3C* | 7.61e-7 |

## Supplementary acknowledgements

We gratefully acknowledge the individual studies and participants who provided biological samples and data for the TOPMed project. Funding for the Barbados Asthma Genetics Study (BAGS) was provided by the National Institutes of Health (NIH) R01HL104608, R01HL087699, and HL104608 S1. The Mount Sinai BioMe Biobank (BioMe) has been supported by The Andrea and Charles Bronfman Philanthropies and in part by funds from the NHLBI and the National Human Genome Research Institute (NHGRI) (U01HG00638001; U01HG007417; X01HL134588); genome sequencing was funded by contract HHSN268201600037I. The Cleveland Clinic Atrial Fibrillation study (CCAF) was supported by NIH grants R01 HL 090620 and R01 HL 111314, the NIH National Center for Research Resources for Case Western Reserve University and Cleveland Clinic Clinical and Translational Science Award UL1-RR024989, the Cleveland Clinic Department of Cardiovascular Medicine philanthropy research funds, and the Tomsich Atrial Fibrillation Research Fund; genome sequencing was supported by R01HL092577. The Framingham Heart Study (FHS) was supported by contracts NO1-HC-25195, HHSN268201500001I, and 75N92019D00031 from the NHLBI and grant supplement R01 HL092577-06S1; genome sequencing was funded by HHSN268201600034I and U54HG003067. The Hypertension Genetic Epidemiology Network Study (HyperGen) is part of the NHLBI Family Blood Pressure Program; collection of the data represented here was supported by grants U01 HL054472, U01 HL054473, U01 HL054495, and U01 HL054509; genome sequencing was funded by R01HL055673. The Jackson Heart Study is supported and conducted in collaboration with Jackson State University (HHSN268201300049C and HHSN268201300050C), Tougaloo College (HHSN268201300048C), and the University of Mississippi Medical Center (HHSN268201300046C and HHSN268201300047C) contracts from NHLBI and the National Institute for Minority Health and Health Disparities (NIMHD); genome sequencing was funded by HHSN268201100037C. The My Life, Our Future samples (MLOF) and data are made possible through the partnership of Blood-works Northwest, the American Thrombosis and Hemostasis Network, the National Hemophilia Foundation, and Bioverativ; genome sequencing was funded by HHSN268201600033I and HHSN268201500016C. The Venous Thromboembolism project (VTE) was funded in part by grants from the NIH, NHLBI (HL66216 and HL83141), and the NHGRI (HG04735). The Vanderbilt Genetic Basis of Atrial Fibrillation study (VUAF) was supported by grants from the American Heart Association (EIA 0940116N) and grants from the National Institutes of Health (HL092217, U19 HL65962, and UL1 RR024975), and by CTSA award (UL1TR000445) from the National Center for Advancing Translational Sciences; genome sequencing was funded by R01HL092577. The Women’s Health Initiative program (WHI) is funded by NHLBI through contracts 75N92021D00001, 75N92021D00002, 75N92021D00003, 75N92021D00004, 75N92021D00005; genome sequencing was funded by HHSN268201500014C.

## References

[1] J. J. Vitti, S. R. Grossman, P. C. Sabeti, Detecting natural selection in genomic data, Annu. Rev. Genet. 47 (2013) 97–120.

[2] V. Pankratov, M. Yunusbaeva, S. Ryakhovsky, M. Zarodniuk, Estonian Biobank Research Team, B. Yunusbayev, Prioritizing autoimmunity risk variants for functional analyses by fine-mapping mutations under natural selection, Nat. Commun. 13 (2022) 7069.

3. A. J. Sams, A. Dumaine, Y. Ńedélec, V. Yotova, C. Alfieri, J. E. Tanner, P. W. Messer, L. B. Barreiro, Adaptively introgressed Neandertal haplotype at the *OAS* locus functionally impacts innate immune responses in humans, Genome Biol. 17 (2016) 246.

4. Anopheles gambiae 1000 Genomes Consortium, Genetic diversity of the African malaria vector Anopheles gambiae, Nature 552 (2017) 96–100.

5. N. R. Garud, Understanding soft sweeps: a signature of rapid adaptation, Nat. Rev. Genet. 24 (2023) 420.

6. B. Guo, V. Borda, R. Laboulaye, M. D. Spring, M. Wojnarski, B. A. Vesely, J. C. Silva, N. C. Waters, T. D. O’Connor, S. Takala-Harrison, Strong positive selection biases identity-by-descent-based inferences of recent demography and population structure in Plasmodium falciparum, Nat. Commun. 15 (2024) 2499.

7. M. Kimura, The Neutral Theory of Molecular Evolution, Cambridge University Press, 1983.

8. T. Ohta, Slightly deleterious mutant substitutions in evolution, Nature 246 (1973) 96–98.

9. J. Hermisson, P. S. Pennings, Soft sweeps: molecular population genetics of adaptation from standing genetic variation, Genetics 169 (2005) 2335–2352.

10. J. Hermisson, P. S. Pennings, Soft sweeps and beyond: understanding the patterns and probabilities of selection footprints under rapid adaptation, Methods Ecol. Evol. 8 (2017) 700–716.

11. P. S. Pennings, J. Hermisson, Soft sweeps III: the signature of positive selection from recurrent mutation, PLoS Genet. 2 (2006) e186.

12. P. S. Pennings, J. Hermisson, Soft sweeps II—molecular population genetics of adaptation from recurrent mutation or migration, Mol. Biol. Evol. 23 (2006) 1076–1084.

13. J. F. Crow, M. Kimura, An Introduction to Population Genetics Theory, Harper & Row, New York, NY, 1970.

14. S. D. Temple, R. K. Waples, S. R. Browning, Modeling recent positive selection using identity-by-descent segments, Am. J. Hum. Genet. 111 (2024) 2510–2529.

15. M. Kreitman, H. Akashi, Molecular evidence for natural selection, Annu. Rev. Ecol. Syst. 26 (1995) 403–422.

16. J. H. McDonald, M. Kreitman, Adaptive protein evolution at the Adh locus in Drosophila, Nature 351 (1991) 652–654.

17. P. C. Sabeti, P. Varilly, B. Fry, J. Lohmueller, E. Hostetter, C. Cotsapas, X. Xie, E. H. Byrne, S. A. McCarroll, R. Gaudet, S. F. Schaffner, E. S. Lander, International HapMap Consortium, Genome-wide detection and characterization of positive selection in human populations, Nature 449 (2007) 913–918.

18. M. Salter-Townshend, S. Myers, Fine-scale inference of ancestry segments without prior knowledge of admixing groups, Genetics 212 (2019) 869–889.

19. J. C. Fay, C. I. Wu, Hitchhiking under positive Darwinian selection, Genetics 155 (2000) 1405–1413.

20. Y. Field, E. A. Boyle, N. Telis, Z. Gao, K. J. Gaulton, D. Golan, L. Yengo, G. Rocheleau, P. Froguel, M. I. McCarthy, J. K. Pritchard, Detection of human adaptation during the past 2000 years, Science 354 (2016) 760.

21. F. Tajima, Statistical method for testing the neutral mutation hypothesis by DNA polymorphism, Genetics 123 (1989) 585–595.

22. A. Akbari, J. J. Vitti, A. Iranmehr, M. Bakhtiari, P. C. Sabeti, S. Mirarab, V. Bafna, Identifying the favored mutation in a positive selective sweep, Nat. Methods 15 (2018) 279–282.

23. A. Albrechtsen, I. Moltke, R. Nielsen, Natural selection and the distribution of identity-by-descent in the human genome, Genetics 186 (2010) 295–308.

24. S. R. Browning, B. L. Browning, Probabilistic estimation of identity by descent segment endpoints and detection of recent selection, Am. J. Hum. Genet. 107 (2020) 895–910.

25. A. Ferrer-Admetlla, M. Liang, T. Korneliussen, R. Nielsen, On detecting incomplete soft or hard selective sweeps using haplotype structure, Mol. Biol. Evol. 31 (2014) 1275–1291.

26. N. R. Garud, P. W. Messer, E. O. Buzbas, D. A. Petrov, Recent selective sweeps in North American Drosophila melanogaster show signatures of soft sweeps, PLoS Genet. 11 (2015) e1005004.

27. A. M. Harris, N. R. Garud, M. DeGiorgio, Detection and classification of hard and soft sweeps from unphased genotypes by multilocus genotype identity, Genetics 210 (2018) 1429–1452.

28. J. Nait Saada, G. Kalantzis, D. Shyr, F. Cooper, M. Robinson, A. Gusev, P. F. Palamara, Identity-by-descent detection across 487,409 British samples reveals fine scale population structure and ultra-rare variant associations, Nat. Commun. 11 (2020) 1–15.

29. P. F. O’Reilly, E. Birney, D. J. Balding, Confounding between recombination and selection, and the Ped/Pop method for detecting selection, Genome Res. 18 (2008) 1304–1313.

30. P. C. Sabeti, D. E. Reich, J. M. Higgins, H. Z. P. Levine, D. J. Richter, S. F. Schaffner, S. B. Gabriel, J. V. Platko, N. J. Patterson, G. J. McDonald, H. C. Ackerman, S. J. Campbell, D. Altshuler, R. Cooper, D. Kwiatkowski, R. Ward, E. S. Lander, Detecting recent positive selection in the human genome from haplotype structure, Nature 419 (2002) 832–837.

31. B. F. Voight, S. Kudaravalli, X. Wen, J. K. Pritchard, A map of recent positive selection in the human genome, PLoS Biol. 4 (2006) e72.

32. P. F. Palamara, J. Terhorst, Y. S. Song, A. L. Price, High-throughput inference of pairwise coalescence times identifies signals of selection and enriched disease heritability, Nat. Genet. 50 (2018) 1311–1317.

33. L. Speidel, M. Forest, S. Shi, S. R. Myers, A method for genome-wide genealogy estimation for thousands of samples, Nat. Genet. 51 (2019) 1321– 1329.

34. A. J. Stern, P. R. Wilton, R. Nielsen, An approximate full-likelihood method for inferring selection and allele frequency trajectories from DNA sequence data, PLoS Genet. 15 (2019) e1008384.

35. A. H. Vaughn, R. Nielsen, Fast and accurate estimation of selection coefficients and allele histories from ancient and modern DNA, Molecular Biology and Evolution 41 (2024) msae156.

36. B. M. Peter, E. Huerta-Sanchez, R. Nielsen, Distinguishing between selective sweeps from standing variation and from a de novo mutation, PLoS Genet. 8 (2012) e1003011.

37. I. Mathieson, J. Terhorst, Direct detection of natural selection in Bronze Age Britain, Genome Res. 32 (2022) 2057–2067.

38. I. Mathieson, I. Lazaridis, N. Rohland, S. Mallick, N. Patterson, S. A. Roodenberg, E. Harney, K. Stewardson, D. Fernandes, M. Novak, K. Sirak, C. Gamba, E. R. Jones, B. Llamas, S. Dryomov, J. Pickrell, J. L. Arsuaga, J. M. B. de Castro, E. Carbonell, F. Gerritsen, A. Khokhlov, P. Kuznetsov, M. Lozano, H. Meller, O. Mochalov, V. Moiseyev, M. A. R. Guerra, J. Rood-enberg, J. M. Vergés, J. Krause, A. Cooper, K. W. Alt, D. Brown, D. Anthony, C. Lalueza-Fox, W. Haak, R. Pinhasi, D. Reich, Genome-wide patterns of selection in 230 ancient Eurasians, Nature 528 (2015) 499–503.

39. H. A. Hejase, Z. Mo, L. Campagna, A. Siepel, A deep-learning approach for inference of selective sweeps from the ancestral recombination graph, Mol. Biol. Evol. 39 (2022).

40. A. D. Kern, D. R. Schrider, diploS/HIC: an updated approach to classifying selective sweeps, G3 (2018).

41. Z. Mo, A. Siepel, Domain-adaptive neural networks improve supervised machine learning based on simulated population genetic data, PLoS Genet. 19 (2023) e1011032.

42. R. Riley, I. Mathieson, S. Mathieson, Interpreting generative adversarial networks to infer natural selection from genetic data, Genetics 226 (2024).

43. D. R. Schrider, A. D. Kern, S/HIC: Robust identification of soft and hard sweeps using machine learning, PLoS Genet. 12 (2016) e1005928.

44. L. S. Whitehouse, D. R. Schrider, Timesweeper: accurately identifying selective sweeps using population genomic time series, Genetics 224 (2023) iyad084.

45. A. J. Stern, R. Nielsen, Detecting natural selection, Handbook of Statistical Genomics: Two Volume Set (2019) 397–340.

46. Z. Sidak, Rectangular confidence regions for the means of multivariate normal distributions, J. Am. Stat. Assoc. 62 (1967) 626–633.

47. Y. Benjamini, Y. Hochberg, Controlling the false discovery rate: a practical and powerful approach to multiple testing, J. R. Stat. Soc. Series B Stat. Methodol. 57 (1995) 289–300.

48. Z. Chen, M. Boehnke, X. Wen, B. Mukherjee, Revisiting the genome-wide significance threshold for common variant GWAS, G3 11 (2021).

49. S. R. Browning, E. A. Thompson, Detecting rare variant associations by identity-by-descent mapping in case-control studies, Genetics 190 (2012) 1521–1531.

50. K. N. Conneely, M. Boehnke, “So many correlated tests, so little time! Rapid adjustment of P values for multiple correlated tests”, Am. J. Hum. Genet. 81 (2007) 1158–1168.

51. K. Grinde, Statistical Inference in Admixed Populations, Ph.D. thesis, University of Washington, 2019.

52. K. E. Grinde, L. A. Brown, A. P. Reiner, T. A. Thornton, S. R. Browning, Genome-wide significance thresholds for admixture mapping studies, Am. J. Hum. Genet. 104 (2019) 454–465.

53. S. R. Seaman, B. Müller-Myhsok, Rapid simulation of P values for product methods and multiple-testing adjustment in association studies, Am. J. Hum. Genet. 76 (2005) 399–408.

54. D. Siegmund, B. Yakir, The Statistics of Gene Mapping, Springer New York, 2007.

55. E. Feingold, P. O. Brown, D. Siegmund, Gaussian models for genetic linkage analysis using complete high-resolution maps of identity by descent, Am. J. Hum. Genet. 53 (1993) 234–251.

56. L. Śegurel, C. Bon, On the evolution of lactase persistence in humans, Annu. Rev. Genomics Hum. Genet. 18 (2017) 297–319.

57. D. Taliun, D. N. Harris, M. D. Kessler, J. Carlson, Z. A. Szpiech, R. Torres, S. A. G. Taliun, A. Corvelo, S. M. Gogarten, H. M. Kang, A. N. Pitsillides, J. LeFaive, S. B. Lee, X. Tian, B. L. Browning, S. Das, A. K. Emde, W. E. Clarke, D. P. Loesch, A. C. Shetty, T. W. Blackwell, A. V. Smith, Q. Wong, X. Liu, NHLBI Trans-Omics for Precision Medicine (TOPMed) Consortium, G. R. Abecasis, Sequencing of 53,831 diverse genomes from the NHLBI TOPMed program, Nature 590 (2021) 290–299.

58. J. M. Granka, B. M. Henn, C. R. Gignoux, J. M. Kidd, C. D. Bustamante, M. W. Feldman, Limited evidence for classic selective sweeps in African populations, Genetics 192 (2012) 1049–1064.

59. S. D. Temple, E. A. Thompson, Identity-by-descent segments in large samples, bioRxiv (2024). doi:10.1101/2024.06.05.597656.

60. S. D. Temple, Statistical Inference Using Identity-by-Descent Segments: Perspectives on Recent Positive Selection, Ph.D. thesis, University of Washington, 2024.

61. K. E. Grinde, B. L. Browning, A. P. Reiner, T. A. Thornton, S. R. Browning, Adjusting for principal components can induce spurious associations in genome-wide association studies in admixed populations, bioRxiv (2024). doi:10.1101/2024.04.02.587682.

62. F. Baumdicker, G. Bisschop, D. Goldstein, G. Gower, A. P. Ragsdale, G. Tsambos, S. Zhu, B. Eldon, E. C. Ellerman, J. G. Galloway, A. L. Gladstein, G. Gorjanc, B. Guo, B. Jeffery, W. W. Kretzschumar, K. Lohse, M. Matschiner, D. Nelson, N. S. Pope, C. D. Quinto-Cortés, M. F. Rodrigues, K. Saunack, T. Sellinger, K. Thornton, H. van Kemenade, A. W. Wohns, Y. Wong, S. Gravel, A. D. Kern, J. Koskela, P. L. Ralph, J. Kelleher, Efficient ancestry and mutation simulation with msprime 1.0, Genetics 220 (2022).

63. S. D. Temple, S. R. Browning, E. A. Thompson, Fast simulation of identity-by-descent segments, bioRxiv (2024). doi:10.1101/2024.12.13.628449.

64. R. Fisher, XXI.—on the dominance ratio, Proceedings of the Royal Society of Edinburgh 42 (1923) 321–341.

65. J. B. S. Haldane, A mathematical theory of natural and artificial selection. Part I, Math. Proc. Camb. Philos. Soc. 23 (1924) 19–41.

66. S. Wright, Evolution in Mendelian populations, Genetics 16 (1931) 97–159.

67. A. D. Kern, D. R. Schrider, Discoal: flexible coalescent simulations with selection, Bioinformatics 32 (2016) 3839–3841.

68. C. Bycroft, C. Freeman, D. Petkova, G. Band, L. T. Elliott, K. Sharp, A. Motyer, D. Vukcevic, O. Delaneau, J. O’Connell, A. Cortes, S. Welsh, A. Young, M. Effingham, G. McVean, S. Leslie, N. Allen, P. Donnelly, J. Marchini, The UK Biobank resource with deep phenotyping and genomic data, Nature 562 (2018) 203–209.

69. B. L. Browning, X. Tian, Y. Zhou, S. R. Browning, Fast two-stage phasing of large-scale sequence data, Am. J. Hum. Genet. 108 (2021) 1880–1890.

70. S. M. Gogarten, T. Sofer, H. Chen, C. Yu, J. A. Brody, T. A. Thornton, K. M. Rice, M. P. Conomos, Genetic association testing using the GENESIS R/Bioconductor package, Bioinformatics 35 (2019) 5346–5348.

71. X. Zheng, D. Levine, J. Shen, S. M. Gogarten, C. Laurie, B. S. Weir, A high-performance computing toolset for relatedness and principal component analysis of SNP data, Bioinformatics 28 (2012) 3326–3328.

72. D. H. Alexander, J. Novembre, K. Lange, Fast model-based estimation of ancestry in unrelated individuals, Genome Res. 19 (2009) 1655–1664.

73. Y. Wu, K. Gettler, M. E. Kars, M. Giri, D. Li, C. S. Bayrak, P. Zhang, A. Jain, P. Maffucci, K. Sabic, T. Van Vleck, G. Nadkarni, L. A. Denson, H. Ostrer, A. P. Levine, E. R. Schiff, A. W. Segal, S. Kugathasan, P. D. Stenson, D. N. Cooper, L. Philip Schumm, S. Snapper, M. J. Daly, T. Haritunians, R. H. Duerr, M. S. Silverberg, J. D. Rioux, S. R. Brant, D. P. B. McGovern, J. H. Cho, Y. Itan, Identifying high-impact variants and genes in exomes of Ashkenazi Jewish inflammatory bowel disease patients, Nat. Commun. 14 (2023) 2256.

74. S. R. Browning, B. L. Browning, Accurate non-parametric estimation of recent effective population size from segments of identity by descent, Am. J. Hum. Genet. 97 (2015) 404–418.

75. S. Carmi, K. Y. Hui, E. Kochav, X. Liu, J. Xue, F. Grady, S. Guha, K. Upadhyay, D. Ben-Avraham, S. Mukherjee, B. M. Bowen, T. Thomas, J. Vijai, M. Cruts, G. Froyen, D. Lambrechts, S. Plaisance, C. Van Broeckhoven, P. Van Damme, H. Van Marck, N. Barzilai, A. Darvasi, K. Offit, S. Bressman, L. J. Ozelius, I. Peter, J. H. Cho, H. Ostrer, G. Atzmon, L. N. Clark, T. Lencz, I. Pe’er, Sequencing an Ashkenazi reference panel supports population-targeted personal genomics and illuminates Jewish and European origins, Nat. Commun. 5 (2014) 4835.

76. X. Tian, B. L. Browning, S. R. Browning, Estimating the genome-wide mutation rate with three-way identity by descent, Am. J. Hum. Genet. 105 (2019) 883–893.

77. A. Raj, M. Stephens, J. K. Pritchard, fastSTRUCTURE: variational inference of population structure in large SNP data sets, Genetics 197 (2014) 573–589.

78. The International HapMap Consortium, The international HapMap project, Nature 426 (2003) 789–796.

79. M. Byrska-Bishop, U. S. Evani, X. Zhao, A. O. Basile, H. J. Abel, A. A. Regier, A. Corvelo, W. E. Clarke, R. Musunuri, K. Nagulapalli, S. Fairley, A. Runnels, L. Winterkorn, E. Lowy, Human Genome Structural Variation Consortium, Paul Flicek, S. Germer, H. Brand, I. M. Hall, M. E. Talkowski, G. Narzisi, M. C. Zody, High-coverage whole-genome sequencing of the expanded 1000 Genomes Project cohort including 602 trios, Cell 185 (2022) 3426–3440.

80. B. V. Halldorsson, G. Palsson, O. A. Stefansson, H. Jonsson, M. T. Hardarson, H. P. Eggertsson, B. Gunnarsson, A. Oddsson, G. H. Halldorsson, F. Zink, S. A. Gudjonsson, M. L. Frigge, G. Thorleifsson, A. Sigurdsson, S. N. Stacey, P. Sulem, G. Masson, A. Helgason, D. F. Gudbjartsson, U. Thorsteinsdottir, K. Stefansson, Characterizing mutagenic effects of recombination through a sequence-level genetic map, Science 363 (2019).

81. R. Cai, B. L. Browning, S. R. Browning, Identity-by-descent-based estimation of the X chromosome effective population size with application to sex-specific demographic history, G3 13 (2023).

82. C. Bhérer, C. L. Campbell, A. Auton, Refined genetic maps reveal sexual dimorphism in human meiotic recombination at multiple scales, Nat. Commun. 8 (2017) 14994.

83. R. M. Gittelman, J. G. Schraiber, B. Vernot, C. Mikacenic, M. M. Wurfel, J. M. Akey, Archaic hominin admixture facilitated adaptation to Out-of-Africa environments, Curr. Biol. 26 (2016) 3375–3382.

84. Y. Horikawa, N. Iwasaki, M. Hara, H. Furuta, Y. Hinokio, B. N. Cockburn, T. Lindner, K. Yamagata, M. Ogata, O. Tomonaga, H. Kuroki, T. Kasahara, Y. Iwamoto, G. I. Bell, Mutation in hepatocyte nuclear factor-1ß gene (*TCF2*) associated with MODY, Nat. Genet. 17 (1997) 384–385.

85. J. Gudmundsson, P. Sulem, V. Steinthorsdottir, J. T. Bergthorsson, G. Thorleifsson, A. Manolescu, T. Rafnar, D. Gudbjartsson, B. A. Agnarsson, A. Baker, et al., Two variants on chromosome 17 confer prostate cancer risk, and the one in *TCF2* protects against type 2 diabetes, Nat. Genet. 39 (2007) 977–983.

86. K. A. Papadakis, J. Prehn, V. Nelson, L. Cheng, S. W. Binder, P. D. Ponath, D. P. Andrew, S. R. Targan, The role of thymus-expressed chemokine and its receptor *CCR9* on lymphocytes in the regional specialization of the mucosal immune system, J. Immunol. 165 (2000) 5069–5076.

87. J. F. Shelton, A. J. Shastri, C. Ye, C. H. Weldon, T. Filshtein-Sonmez, D. Coker, A. Symons, J. Esparza-Gordillo, 23andMe COVID-19 Team, S. Aslibekyan, A. Auton, Trans-ancestry analysis reveals genetic and nongenetic associations with COVID-19 susceptibility and severity, Nat. Genet. 53 (2021) 801–808.

88. S. R. Browning, B. L. Browning, Y. Zhou, S. Tucci, J. M. Akey, Analysis of human sequence data reveals two pulses of archaic Denisovan admixture, Cell 173 (2018) 53–61.

89. Q. Ding, Y. Hu, S. Xu, J. Wang, L. Jin, Neanderthal introgression at chromosome 3p21.31 was under positive natural selection in East Asians, Mol. Biol. Evol. 31 (2014) 683–695.

90. H. Stefansson, A. Helgason, G. Thorleifsson, V. Steinthorsdottir, G. Masson, J. Barnard, A. Baker, A. Jonasdottir, A. Ingason, V. G. Gudnadottir, et al., A common inversion under selection in Europeans, Nat. Genet. 37 (2005) 129–137.

91. I. G. Romero, C. Basu Mallick, A. Liebert, F. Crivellaro, G. Chaubey, Y. Itan, M. Metspalu, M. Eaaswarkhanth, R. Pitchappan, R. Villems, D. Reich, L. Singh, K. Thangaraj, M. G. Thomas, D. M. Swallow, M. Mirazón Lahr, T. Kivisild, Herders of Indian and European cattle share their predominant allele for lactase persistence, Mol. Biol. Evol. 29 (2012) 249– 260.

92. K. K. Kidd, A. J. Pakstis, M. P. Donnelly, O. Bulbul, L. Cherni, C. Gurkan, L. Kang, H. Li, L. Yun, P. Paschou, K. A. Meiklejohn, E. Haigh, W. C. Speed, The distinctive geographic patterns of common pigmentation variants at the *OCA2* gene, Sci. Rep. 10 (2020) 15433.

93. E. Oancea, J. Vriens, S. Brauchi, J. Jun, I. Splawski, D. E. Clapham, *TRPM1* forms ion channels associated with melanin content in melanocytes, Science Signaling 2 (2009) ra21–ra21.

94. M. Lewis, S. Vyse, A. Shields, S. Boeltz, P. Gordon, T. Spector, P. Lehner, H. Walczak, T. Vyse, *UBE2L3* polymorphism amplifies NF-*κ*B activation and promotes plasma cell development, linking linear ubiquitination to multiple autoimmune diseases, Am. J. Hum. Genet. 96 (2015) 221–234.

95. J. Schreml, B. Durmaz, O. Cogulu, K. Keupp, F. Beleggia, E. Pohl, E. Milz, M. Coker, S. K. Ucar, G. Nürnberg, P. Nürnberg, J. Kuhn, F. Ozkinay, The missing “link”: an autosomal recessive short stature syndrome caused by a hypofunctional *XYLT1* mutation, Hum. Genet. 133 (2014) 29–39.

96. E. K. Mis, K. F. Liem, Jr, Y. Kong, N. B. Schwartz, M. Domowicz, S. D. Weatherbee, Forward genetics defines *XYLT1* as a key, conserved regulator of early chondrocyte maturation and skeletal length, Dev. Biol. 385 (2014) 67–82.

97. S. F. Oster, M. O. Bodeker, F. He, D. W. Sretavan, Invariant *SEMA5A* inhibition serves an ensheathing function during optic nerve development, Development 130 (2003) 775–784.

98. A.-L. Mosca-Boidron, L. Gueneau, G. Huguet, A. Goldenberg, C. Henry, N. Gigot, E. Pallesi-Pocachard, A. Falace, L. Duplomb, J. Thevenon, Y. Duffourd, J. St-Onge, P. Chambon, J.-B. Rivìere, C. Thauvin-Robinet, P. Callier, N. Marle, M. Payet, C. Ragon, H. Goubran Botros, J. Buratti, S. Calderari, G. Dumas, R. Delorme, N. Lagarde, J.-M. Pinoit, A. Rosier, A. Masurel-Paulet, C. Cardoso, F. Mugneret, P. Saugier-Veber, D. Campion, L. Faivre, T. Bourgeron, A de novo microdeletion of *SEMA5A* in a boy with autism spectrum disorder and intellectual disability, Eur. J. Hum. Genet. 24 (2016) 838–843.

99. J. Hao, X. Han, H. Huang, X. Yu, J. Fang, J. Zhao, R. A. Prayson, S. Bao, J. S. Yu, *SEMA3C* signaling is an alternative activator of the canonical WNT pathway in glioblastoma, Nat. Commun. 14 (2023) 2262.

100. R. C. Hardison, Evolution of hemoglobin and its genes, Cold Spring Harb. Perspect. Med. 2 (2012) a011627.

101. S. A. Tishkoff, S. M. Williams, Genetic analysis of African populations: human evolution and complex disease, Nat. Rev. Genet. 3 (2002) 611–621.

102. A. Freudiger, V. M. Jovanovic, Y. Huang, N. Snyder-Mackler, D. F. Conrad, B. Miller, M. J. Montague, H. Westphal, P. F. Stadler, S. Bley, J. E. Horvath, L. J. N. Brent, M. L. Platt, A. Ruiz-Lambides, J. Tung, K. Nowick, H. Ringbauer, A. Widdig, Estimating realized relatedness in free-ranging macaques by inferring identity-by-descent segments, Proceedings of the National Academy of Sciences 122 (2025) e2401106122.

103. H. Ringbauer, Y. Huang, A. Akbari, S. Mallick, I. Olalde, N. Patterson, D. Reich, Accurate detection of identity-by-descent segments in human ancient dna, Nat. Genet. 56 (2024) 143–151.

104. I. V. Caldas, A. G. Clark, P. W. Messer, Inference of selective sweep parameters through supervised learning, bioRxiv (2022). doi:10.1101/2022.07.19.500702.

105. L. S. Whitehouse, D. D. Ray, D. R. Schrider, Tree sequences as a general-purpose tool for population genetic inference, Mol. Bio. Evol. 41 (2024) msae223.

106. M. Hernandez, G. H. Perry, Scanning the human genome for “signatures” of positive selection: transformative opportunities and ethical obligations, Evol. Anthropol. 30 (2021) 113–121.

107. A. Akbari, A. R. Barton, S. Gazal, Z. Li, M. Kariminejad, A. Perry, Y. Zeng, A. Mittnik, N. Patterson, M. Mah, et al., Pervasive findings of directional selection realize the promise of ancient DNA to elucidate human adaptation, bioRxiv (2024). doi:10.1101/2024.09.14.613021.

108. C. E. G. Amorim, K. Nunes, D. Meyer, D. Comas, M. C. Bortolini, F. M. Salzano, T. Hünemeier, Genetic signature of natural selection in first Americans, Proceedings of the National Academy of Sciences 114 (2017) 2195– 2199.

109. J. Bryk, E. Hardouin, I. Pugach, D. Hughes, R. Strotmann, M. Stoneking, S. Myles, Positive selection in East Asians for an *EDAR* allele that enhances NF-*κ*b activation, PloS one 3 (2008) e2209.

110. L. Skov, M. C. Macia, E. A. Lucotte, M. I. A. Cavassim, D. Castellano, M. H. Schierup, K. Munch, Extraordinary selection on the human X chromosome associated with archaic admixture, Cell Genomics 3 (2023).

111. A. R. V. R. Horimoto, D. Xue, T. A. Thornton, E. E. Blue, Admixture mapping reveals the association between Native American ancestry at 3q13.11 and reduced risk of Alzheimer’s disease in Caribbean Hispanics, Alzheimers. Res. Ther. 13 (2021) 122.

112. A. Jiménez-Kaufmann, A. Y. Chong, A. Cortés, C. D. Quinto-Cortés, S. L. Fernandez-Valverde, L. Ferreyra-Reyes, L. P. Cruz-Hervert, S. G. Medina-Muñoz, M. Sohail, M. J. Palma-Martinez, G. Delgado-Sánchez, N. Mongua-Rodŕıguez, A. J. Mentzer, A. V. S. Hill, H. Moreno-Maćıas, A. Huerta-Chagoya, C. A. Aguilar-Salinas, M. Torres, H. L. Kim, N. Kalsi, S. C. Schuster, T. Tusíe-Luna, D. O. Del-Vecchyo, L. Garćıa-Garćıa, A. Moreno-Estrada, Imputation performance in Latin American populations: Improving rare variants representation with the inclusion of Native American genomes, Front. Genet. 12 (2021) 719791.

113. R. A. Maller, G. Müller, A. Szimayer, Ornstein–Uhlenbeck processes and extensions, Handbook of Financial Time Series (2009) 421–437.

